# Vutiglabridin improves neurodegeneration in MPTP-induced Parkinson’s disease mice by targeting mitochondrial paraoxonase-2

**DOI:** 10.1101/2022.10.20.512990

**Authors:** Sora Kang, Leo S. Choi, Suyeol Im, Ji Hwan Kim, Keun Woo Lee, Dong Hwan Kim, Jung Hee Park, Min-Ho Park, Jaemin Lee, Sun Kyung Park, Kwang Pyo Kim, Hyeong Min Lee, Hyun Ju Jeon, Hyung Soon Park, Sang-Ku Yoo, Youngmi Kim Pak

## Abstract

Parkinson’s disease (PD), characterized by degeneration of dopaminergic neurons, share pathogenic features with obesity, including mitochondrial dysfunction and oxidative stress. Paraoxonase 2 (PON2) is an inner mitochondrial membrane protein that is highly expressed in dopaminergic neurons and is involved in the regulation of mitochondrial oxidative stress. However, no drug targeting PON2 has ever been developed for the treatment of PD. Here, we show that vutiglabridin, a clinical phase 2-stage drug for the treatment of obesity, has therapeutic effects in PD models, targeting mitochondrial PON2. Vutiglabridin penetrates into the brain, binds to PON2, and restores 1-methyl-4-phenylpyridinium (MPP^+^)-induced mitochondrial dysfunction in SH-SY5Y neuroblastoma cells. Knockdown of PON2 by lentiviral shRNA infection abolished the effects of vutiglabridin on mitochondria. In mice, vutiglabridin significantly alleviated motor impairments and damage to dopaminergic neurons in 1-methyl-4-phenyl-1,2,3,6-tetrahydropyridine (MPTP)-induced PD model, and these effects were also abolished in PON2-knockdown mice, suggesting that vutiglabridin is neuroprotective via PON2. Extensive in vitro and in vivo assessment of potential neurotoxicity showed vutiglabridin to be safe. Overall, these findings provide support for the clinical development of vutiglabridin as a novel PON2 modulator for the treatment of PD.

**One Sentence Summary:** Targeting paraoxonase-2 by a clinical-stage compound vutiglabridin provides neuroprotective effects in preclinical models of Parkinson’s disease.

## INTRODUCTION

Parkinson’s disease (PD) is the second most common and one of the world’s fastest growing neurodegenerative disease (*1, 2*). PD is characterized by the selective loss of tyrosine hydroxylase (TH)-positive dopaminergic neurons in substantia nigra parts compacta (SNpc) (*3*). Evidence strongly suggests that mitochondrial dysfunction and oxidative stress are involved in its pathogenesis (*4, 5*). The main mitochondrial defect in PD is associated with complex I (NADH-ubiquinone oxidoreduction) of the mitochondrial oxidative phosphorylation (OXPHOS); complex I activity was found to be decreased in post-mortem SNpc (*6, 7*), and PD-inducing environmental toxin 1-methyl-4-phenyl-1,2,3,6-tetrahydropyridine (MPTP) is reported to inhibit complex I with its active metabolite 1-methyl-4-phenylpyridinium ion (MPP^+^) (*8, 9*). Also, dopaminergic neurons are particularly prone to oxidative stress as the presence of both dopamine being autoxidized into quinone form and iron that catalyzes Fenton reaction generates high amounts of superoxide radicals and hydrogen peroxide (*10, 11*). Mitochondrial dysfunction observed in PD patients further increases the generation of reactive oxygen species (ROS) including superoxides, and consequent oxidative stress can trigger a cascade of events that lead to the death of dopaminergic neurons (*12*).

Paraoxonase 2 (PON2) is a ubiquitously expressed intracellular membrane protein that has a lactonase enzyme activity and an antioxidant property of specifically reducing mitochondrial superoxide release (*13–15*). As expected from a phylogenetic analysis of the paraoxonase gene family (PON1, PON2, and PON3) showing that PON1 and PON3 evolved from PON2, (*16*), PON2 shares about 70% sequence identity to PON1 (*17*). PON1 has been extensively studied as a plasma HDL-associated protein that reduces lipid peroxidation and is strongly linked to atherosclerosis (*18*), and PON2 is similarly known to have anti-atherogenic effect (*19*). Among the PON family proteins, PON2 is unique among its family to be expressed in the brain tissue, in both rodents and human (*20*). In brain, PON2 expression levels are highest in the dopaminergic regions – SNpc, striatum (ST), and nucleus accumbens – compared to other areas, and PON2 is primarily localized to mitochondria (*21*). PON2 has been reported to specifically reduce mitochondrial ROS level; it binds to coenzyme Q_10_, a cofactor for the electron transport to mitochondrial complex III, and its deficiency in mice increases mitochondrial superoxide production (*22*). Because of its distinct distribution to dopaminergic neurons and its mitochondria-specific anti-oxidative property, PON2 has been speculated to provide neuroprotection against PD (*21–24*). However, there are currently no therapeutic agents being developed that target PON2 for the treatment of PD.

Vutiglabridin (Vuti, HSG4112, **Figure 1A**) is a synthetic new chemical that is currently in human phase 2 clinical study (NCT05197556) for the treatment of obesity (*25*). Vutiglabridin is a derivative of glabridin (*25*), a natural prenylated polyphenolic isoflavan and an important constituent of licorice root (*Glycyrrhiza glabra*), and it is a racemic mixture of (R)-vutiglabridin and (S)-vutiglabridin. Glabridin is associated with numerous biological properties ranging from antioxidant, anti-inflammatory (*26, 27*), neuroprotective (*28–30*), and anti-atherogenic effects to the regulation of energy metabolism (*31*). It also ameliorates lipid dysregulation, and activates the signaling pathway for AMP-protein kinase (AMPK) (*32*), an enzyme well-known for sensing low intracellular ATP levels and controlling mitochondrial biogenesis (*33*). Since mitochondrial dysfunction is suggested as a common pathophysiological mechanism of obesity and neurodegenerative diseases, glabridin is a good candidate for treating both obesity and neurodegenerative diseases. However, because glabridin has several limitations as a therapeutic medicine, such as low physicochemcal stability, low bioavailability, and very low blood-brain barrier (BBB) permeability (*26, 34*), vutiglabridin was designed to improve its druggability.

**Fig 1.**
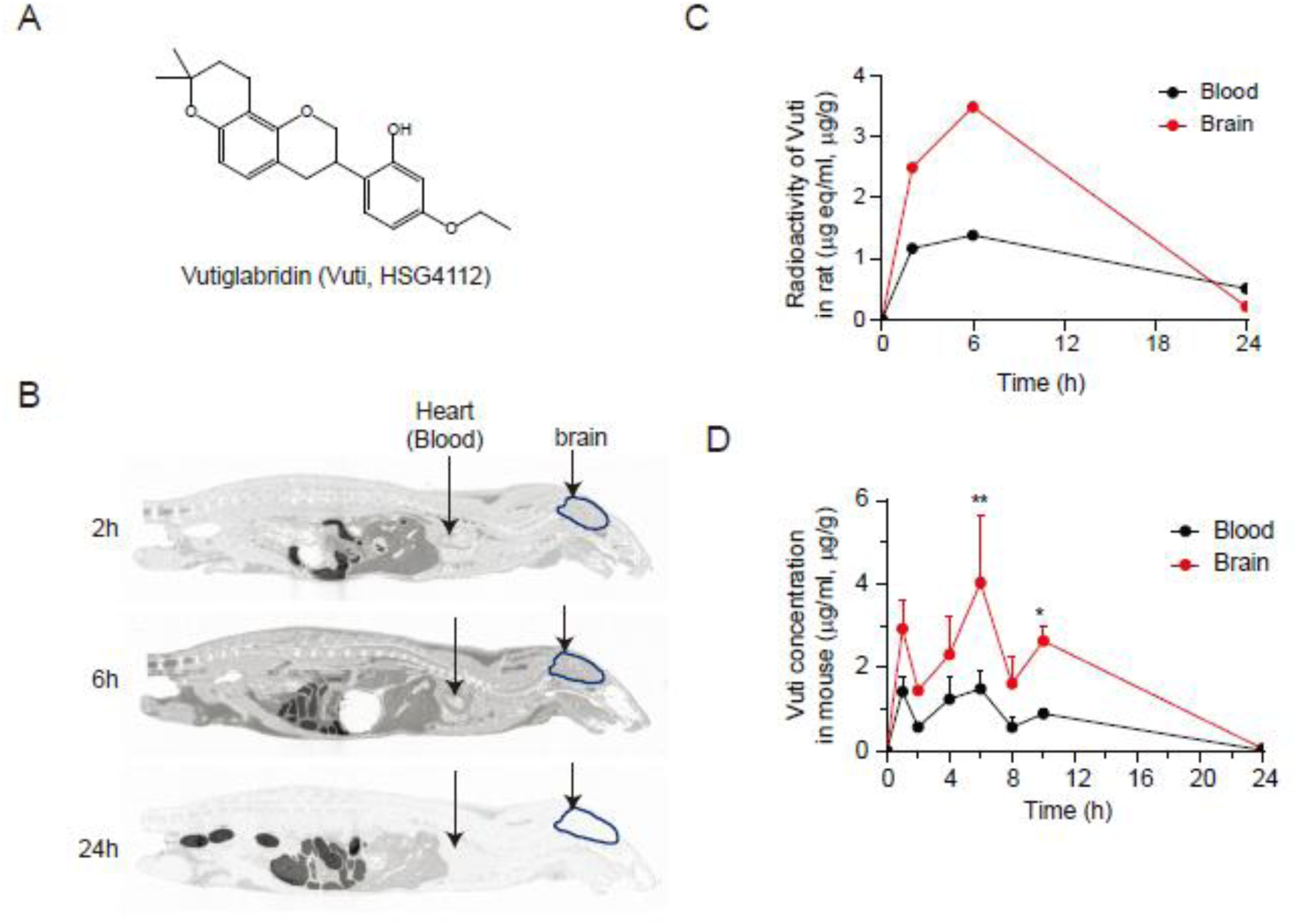
Vutiglabridin penetrates into the brain. (A) Chemical structure of vutiglabridin. Asterisk denotes stereocenter. (B) Whole-body autoradiograms shown from central axis aspect at 2 h, 6 h, and 24 h after a single oral administration of 10 mg/kg of ^14^C-labeled vutiglabridin to a fasted male SD rat. Arrows show the site of brain and heart (blood). (C) Comparative radioactivity of ^14^C-labeled vutiglabridin (Vuti) in the brain and blood (n = 1 per group per timepoint). (D) Concentration of vutiglabridin in the plasma and brain of male C57BL/6J mice after a single oral administration of 50 mg/kg measured via LC-MS/MS analysis. The data are plotted as mean ± SEM (n = 3 per timepoints at 0, 1, 2, 4, 6, 8, 10, and 24 h). \**p* < 0.05, \*\**p* < 0.01 vs. Blood. *p* values are from a two-way ANOVA followed by Sidak’s test.

It is interesting to note several reports showing that glabridin is associated with PON2. A direct interaction between glabridin and PON2 was shown by fluorescence quenching study of glabridin-PON2 in vitro (*35*). Glabridin increased the enzyme activity and the expression of PON2 in THP-1 human monocyte under high glucose stress, as well as in heart and liver tissues of hyperglycemic mice (*35, 36*). Therefore, we focused on PON2 as a target molecule linking glabridin with mitochondria in brain.

In the present study, we investigated whether vutiglabridin modulates mitochondrial activity, and interacts with PON2 to show any neuroprotective effects in PD models. We found that vutiglabridin directly binds to PON2 and is rapidly distributed in the brain. Vutiglabridin restored MPP^+^-induced damages to mitochondria in neuronal cells and significantly alleviated dopaminergic cell death and motor behavior in MPTP-injected PD mice. PON2 knock-down abrogated all beneficial effects of vutiglabridin in PD models, demonstrating that PON2 is a molecular target of vutiglabridin. Our results suggest that vutiglabridin is a promising therapeutic candidate for PD and that PON2 is a novel therapeutic target for PD treatment.

## RESULTS

### Vutiglabridin penetrates the blood-brain barrier in rodents

**Figure 1A** shows the chemical structure of vutiglabridin, which is a very hydrophobic compound with octanol-water partition coefficient logD > 3.69 (**Supplemental Figure S1A**). Given its hydrophobicity and small molecular weight (354.44 g/mol), vutiglabridin is highly likely to penetrate BBB. To examine whether vutiglabridin is permeable across the BBB, an in vitro BBB permeability test was performed by evaluating whether vutiglabridin is a substrate of P-glycoprotein (permeability glycoprotein, P-gp or multidrug resistance protein 1, MDR1), which is an ATP-dependent efflux transporter expressed in the brain endothelial cells that effluxes many hydrophobic compounds out of the cell (*37*). The basal and apical concentration of vutiglabridin was measured by LC-MS/MS analysis in a standard set of cell monolayers: MDCKII-hMDR1, MDCKII, and Caco-2 cell monolayers. Evaluating the efflux ratio with or without P-gp inhibitor showed that vutiglabridin has very low efflux ratio (≤ 2) in all conditions, indicating that vutiglabridin is not a P-gp substrate (Supplementary **Figure S1B**).

Next, the distribution of vutiglabridin to the brain was monitored by a standard quantitative whole-body autoradiography (QWBA) study in Sprague-Dawley (SD) rat (*38*) using ^14^C-labeled vutiglabridin (**Supplemental Figure S1C** for synthesis reaction). Single oral administration of 10 mg/kg of ^14^C-labeled vutiglabridin reached maximum plasma concentration at 6 h (T_max_) (**Supplemental Figure S1D**). In order to quantify spatial and temporal distribution of ^14^C-labeled vutiglabridin, whole-body sagittal sections were made on a different set of SD rats that were individually sacrificed at 2 h, 6 h, and 24 h timepoint. ^14^C-labeled vutiglabridin was distributed throughout most organs in the body except for bone, eyeball, and seminal vesicle (**Figure 1B, Supplemental Table S1**). When radioactivity densities of the image of sagittal sections were quantified based on the blood and brain area, ^14^C-labeled vutiglabridin was successfully distributed to the brain at a level 2.5-fold compared to the plasma at T_max_ of 6 h (**Figure 1C**).

Brain penetration by vutiglabridin was further evaluated in C57BL/6J mice. Single oral administration of 50 mg/kg of vutiglabridin showed that the drug concentration in the brain was 2.7-fold greater than plasma at T_max_ of 6 h (4.03 μg/g in brain vs. 1.49 μg/ml in plasma) and was 2.6-fold greater than plasma in the overall area under the curve exposure (AUC_24h_) (42.5 μg.h/g in brain vs. 16.1 μg.h/ml in plasma) (**Figure 1D**). These results consistently show that vutiglabridin penetrates the BBB and is highly distributed to the brain.

### Vutiglabridin protects against MPP^+^-induced cellular and mitochondrial damage in neuronal cells

We evaluated whether vutiglabridin mediates protective effect against mitochondrial damage by modulating PON2 under the condition of using MPP^+^ toxin as a mitochondrial complex I inhibitor as a model of PD. We first treated 1 mM of MPP^+^ to SH-SY5Y human neuronal cells for 24 h to induce mitochondrial and cellular damage and then post-treated vutiglabridin for 24 h. Vutiglabridin dose-dependently and significantly restored MPP^+^-induced mitochondrial and cellular damage as evidenced by restoring NADH dehydrogenase complex 1 activity (methyl thiazyl tetrazolium; MTT), intracellular ATP content, cellular ROS (DCF-DA), and mitochondrial superoxide (MitoSOX) (**Figure 2A-D**). We also measured oxygen consumption rate (OCR), which is an essential physiological indicator of cellular mitochondrial function (*39*), and found that vutiglabridin significantly rescues MPP^+^-induced reduction in basal respiration, ATP turnover rate, maximum respiratory capacity, and ATP production rate (**Figure 2E-I**). These data demonstrate that vutiglabridin is remarkably effective at ameliorating the neurotoxin-induced mitochondrial dysfunction and cellular damage in neuronal cells.

**Fig 2.**
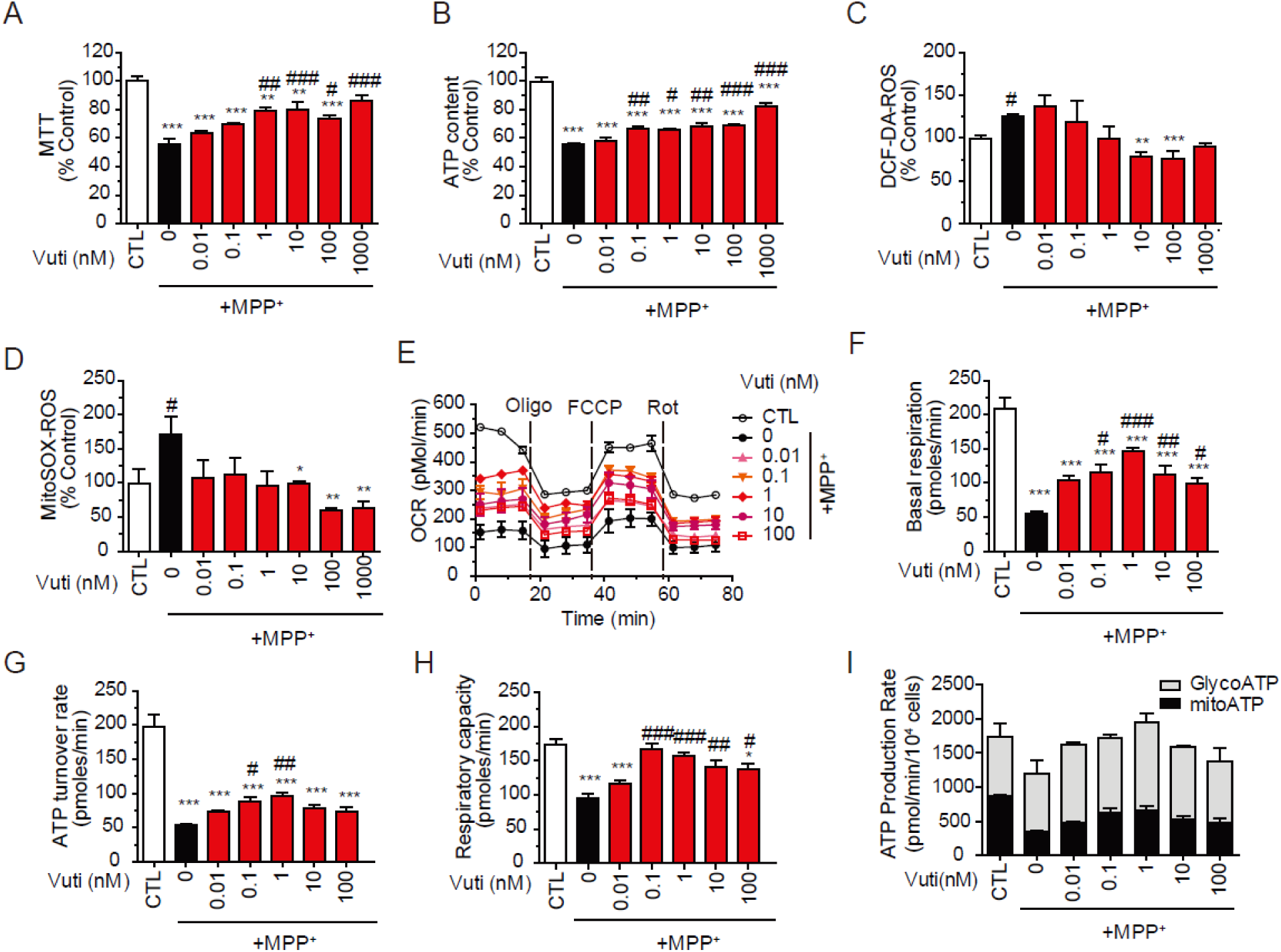
Dose-dependent effects of vutiglabridin on MPP^+^-induced cellular and mitochondrial damage in SH-SY5Y neuronal cells. SH-SY5Y cells were placed in a 96-well plate and incubated with 1 mM MPP^+^ for 24 h, and then were post-treated with vutiglabridin (Vuti, 0-1 µM) for 24 h. The mitochondrial activity was analyzed. (A) MTT as NADH dehydrogenase activity of OXPHOS complex 1. (B) Intracellular ATP content. (C) DCF-DA-based total reactive oxygen species (ROS) generation. (D) MitoSox-based mitochondrial superoxide generation. (E-G) SH-SY5Y cells were placed in an XF-24 microplate and incubated with 1mM MPP^+^ for 20h, and then were post-treated with vutiglabridin (Vuti, 1-100 nM) for 24 h. The oxygen consumption rates (OCR) were analyzed using a Seahorse XF-24 analyzer. (E) OCR profile. Oligomycin (Oligo), FCCP, and rotenone (Rot) were consecutively injected to obtain mitochondrial respiratory capacities. (F) Basal respiration OCR. (G) ATP turnover rate (basal OCR – oligomycin-inhibited OCR). (H) Total respiratory capacity (FCCP-induced OCR). (I) Using ECAR and OCR profiles, ATP production rates through glycolysis (GlycoATP, grey bar) or mitochondria (mitoATP, black bar) were calculated. Mitochondrial activity values are reported as a percentage of the control (CTL). The data are plotted as the mean ± SEM (n = 3). \**p* < 0.05, \*\**p* < 0.01, \*\*\**p* < 0.001 vs. CTL (white bar); ^#^*p* < 0.05, ^##^*p* < 0.01, ^###^*p* < 0.001 vs. MPP^+^-treated control (Black bar). *p* values are from a one-way ANOVA followed by Tukey’s test.

### Vutiglabridin protects against damage on dopaminergic neuron and motor behavior in MPTP-injected mice

Next, we examined whether the in vitro therapeutic effects of vutiglabridin against neurotoxin is translated to in vivo models of PD. MPTP, which is injected intraperitoneally and passes the BBB to be converted by the enzyme monoamine oxidase B (MAO-B) into the neurotoxin MPP^+^, was used to sub-acutely induce PD in C57BL/6J mice. The experimental scheme is shown in **Figure 3A**. Briefly, a day before sacrifice, rotarod and pole test were performed to assess motor behavioral changes, and at sacrifice, mouse brains were collected. Brain sections were immunostained with anti-TH antibody and dopaminergic neuronal damage was analyzed. Vutiglabridin at 0, 1, 50 and 100 mg/kg/day were administered for three weeks, while 0.1 mg/kg rasagiline, an irreversible MAO-B inhibitor that prevents MPTP conversion into the toxic MPP^+^, was administered as a positive reference drug. MPTP-injected mice showed significant reduction in latency to fall in rotarod test and latency time to arrive at the floor (T-LA) in the pole test, as well as significant depletion of TH-positive cells in the SNpc and loss of TH-positive fibers in the ST, compared to the vehicle-only administered (Normal) group (**Figure 3B-F**). Three-week treatment of both vutiglabridin and rasagiline significantly increased latency time in the rotarod test and decreased T-LA in the pole test and increased the number of TH-positive neurons in SNpc and fiber in ST (**Figure 3B-F**). These findings suggest that vutiglabridin, consistent with the findings in neuronal cells, indeed ameliorates neurotoxin-induced damage in PD mice. No clear dose-dependent increase in the neuroprotective effect was observed between 50 and 100 mg/kg of vutiglabridin, so 50 mg/kg was set as the maximum efficacy dose.

**Fig 3.**
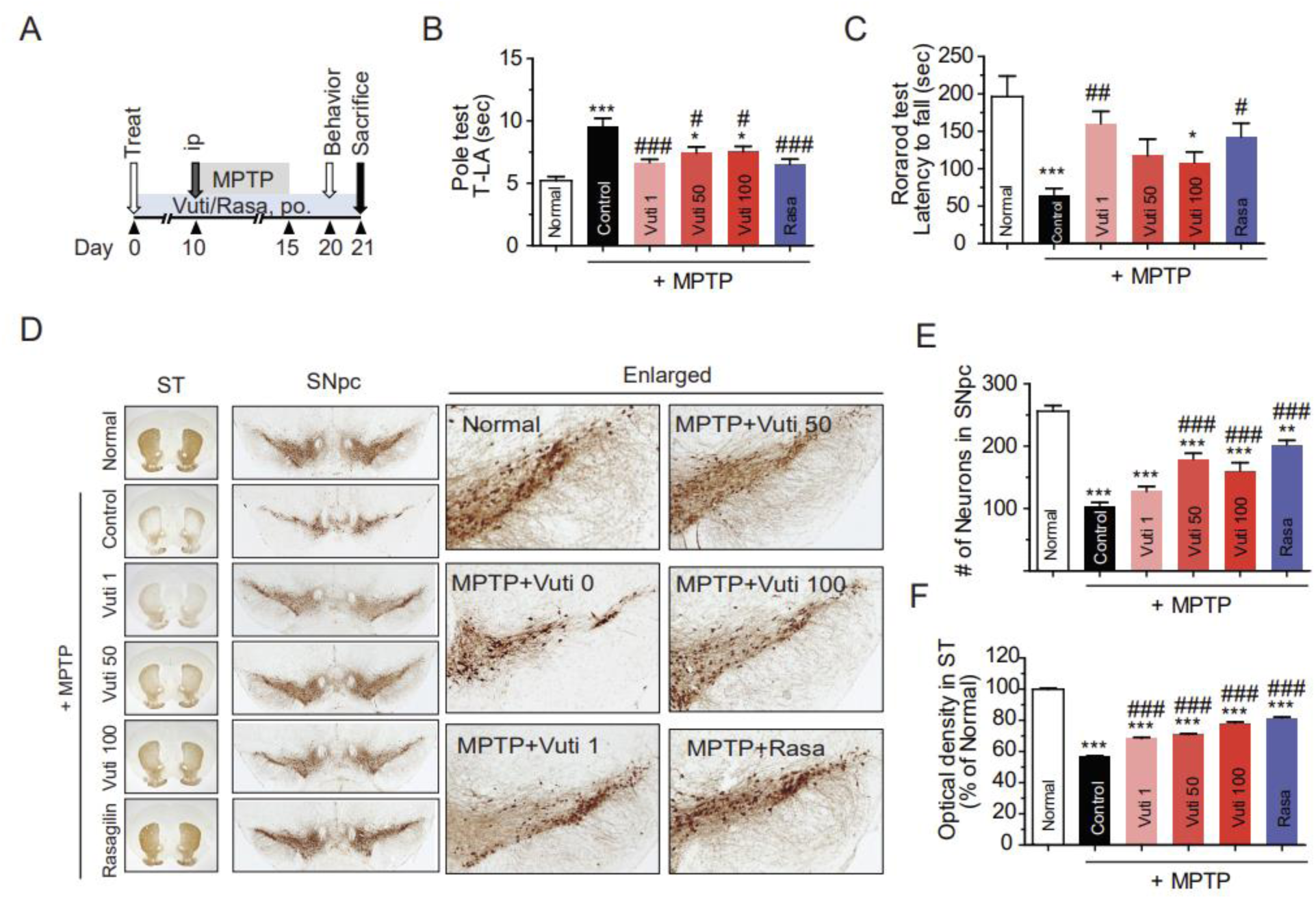
Vutiglabridin attenuates motor impairment and protects dopaminergic neurons in MPTP-injected PD mice. (A) Scheme of the experimental design. C57BL/6 mice (male 8-10 weeks old, n=5 per group) were orally administered with vutiglabridin (Vuti, 1, 50 or 100 mg/kg/day) or rasagiline (Rasa, 0.1 mg/kg/day) for 21 consecutive days. All groups except for the normal control group were injected with MPTP intraperitoneally at 30 mg/kg/day for five days starting on day 10. On day 20, the rotarod and pole test were performed. On day 21, mice were sacrificed, and dopaminergic neurons were visualized via tyrosine hydroxylase (TH) immunohistochemistry. (B) Latency time in the rotarod test. (C) Latency time to arrive at the floor (T-LA) in the pole test. (D) Representative photomicrographs of the SNpc and the striatum (ST). (E) Stereological count of the number of TH-immunopositive neurons in SNpc. (F) Optical density in the ST. The data are plotted as the mean ± SEM (n = 5). \**p* < 0.05, \*\**p* < 0.01, \*\*\**p* < 0.001 vs. vehicle-only group; ^#^*p* < 0.05, ^###^*p* < 0.001 vs. vehicle-only group injected with MPTP. *p* values are from a one-way ANOVA followed by Tukey’s test.

### Vutiglabridin binds to PON2 and increases its protein stability

In order to assess potential binding interaction of vutiglabridin to PON2, we constructed a de novo three-dimensional (3D) in silico model of PON2 based on a homology modeling to its paraoxonase family member PON1, which shares 61.7% sequence identity and 79.2% sequence similarity (**Supplemental Figure S2**). This was performed because the full protein structure of PON2 is unknown, while that of PON1 is known. The structural stability of the constructed PON2 was verified through molecular dynamic simulation, which showed stable maintenance for 10 ns and adequate interatomic collisions as viewed by the Ramachandran plot (**Supplemental Figure S2**). Molecular docking simulation showed that both (R) and (S) form of vutiglabridin showed hydrophobic and non-hydrophobic (polar and electrostatic) interactions with PON2 (**Figure 4A-D**); they had cluster match of 44 and 38 out of 50, and Genetic Optimization for Ligan Docking (GOLD) fitness score of 60.4 and 58.5, respectively. Glabridin also had similar binding interactions with PON2, with lower scores in cluster match and in GOLD fitness score, indicating that vutiglabridin binds to PON2 better than glabridin.

**Fig 4.**
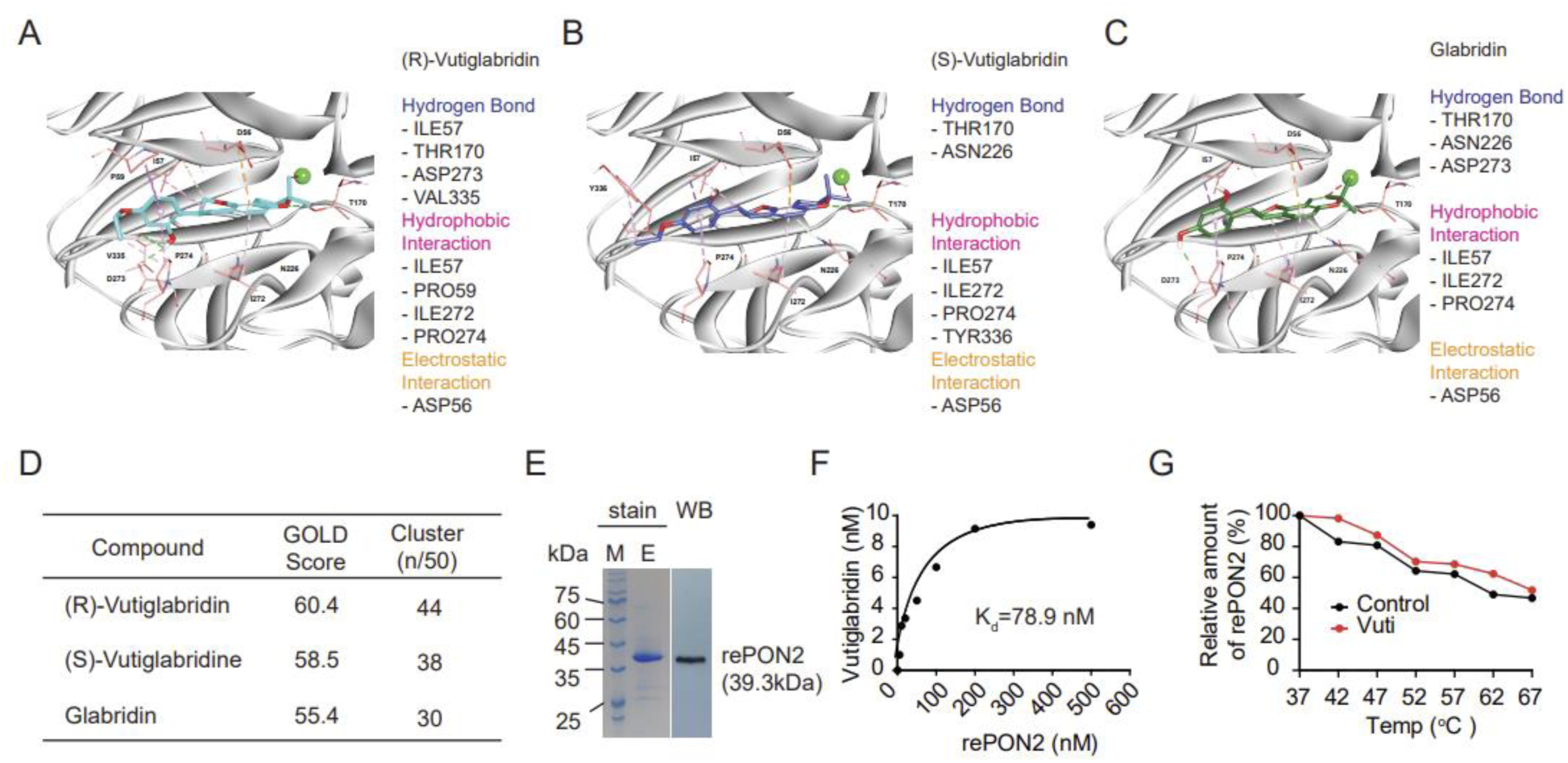
De novo 3D structure of PON2 and binding interaction with vutiglabridin. (A-C) Binding site of PON2 and the molecular docking simulation of (R)-Vutiglabridin (A), (S)-Vutiglabridin, (B), and glabridin (C) to PON2, showing sites of hydrogen bonding, hydrophobic interaction, and electrostatic interaction. (D) Scoring of molecular docking by GOLD fitness score and cluster match analysis. (E) Coomassie blue staining of SDS-PAGE and Western blot of final purified recombinant PON2 (rePON2). (F) RePON2 at 0, 5, 10, 20, 50, 100, 200, and 500 nM was incubated with 10 nM of vutiglabridin in tube for 30 min, and PON2-bound vutiglabridin was isolated and analyzed in quantity by LC-MS/MS. Dissociation constant (K_d_) was 78 nM. The data are plotted as the mean ± SEM (n = 3; error bar is not visible because the error range is negligible). (G) Vutiglabridin (10 nM) was incubated with 500 nM rePON2 for 30 min under 37-67 °C and the relative protein abundance of non-denatured form of rePON2 was measured via LC-MS/MS.

To further confirm vutiglabridin binding to PON2, a mass spectrometry-based binding assay (*40, 41*) was performed using a stable and soluble recombinant human PON2 (rePON2) with C-terminal His-tag, which was newly created for this study (**Figure 4E, Supplemental Materials**) (*42–44*). In tube, various concentrations of rePON2 were incubated with 10 nM of vutiglabridin, and the amount of rePON2-bound vutiglabridin was analyzed using LC-MS/MS. PON2-bound vutiglabridin dose-dependently increased, and the dissociation constant (K_d_) was calculated to be 78 nM (**Figure 4F**), which show that vutiglabridin binds to PON2 with relatively high affinity in the nanomolar range.

To examine how the binding of vutiglabridin affects PON2, the PON2 protein stability was measured via thermal shift assay, which measures the change in thermal denaturation. Here, 500 nM rePON2 was incubated with 10 nM of protein under varying temperatures (37-67 °C) for 30 min, and non-denatured form of PON2 was measured by LC-MS/MS analysis. Vutiglabridin increased the relative protein abundance of non-denatured form of rePON2 at all temperatures (**Figure 4G**), which suggest that vutiglabridin enhances the protein stability of PON2.

### Knockdown of PON2 abolishes vutiglabridin-mediated effects on MPP+-treated neuronal cells

Next, we investigated whether PON2 is indeed required for the neuroprotective effects of vutiglabridin against MPP^+^-induced damage in neuronal cells. We established a PON2 knockdown (KD) model by infecting SH-SY5Y cells with lentiviral particles of PON2 shRNA (shPON2). Western blot and real-time qPCR analyses revealed that approximately 60% PON2 protein expression (both glycosylated and non-glycosylated PON2) is reduced (**Figure 5A, B**) in shPON2-infected PON2-KD (shPON2) cells. In control cells infected with control shRNA (shSCR), vutiglabridin significantly restored MPP^+^-induced mitochondrial and cellular damage similarly to Figure 2. However, in shPON2 cells, the effects of vutiglabridin on mitochondria function were significantly and almost completely abrogated, as measured by MTT, ATP content, TMRE, DCF-DA, and MitoSOX (**Figure 5C-G**). In addition, measurement of OCR showed that while vutiglabridin significantly restored MPP^+^-induced damage in basal respiration, ATP turnover rate, and maximum respiratory capacity in control shSCR cells, and such effects were not observed in shPON2-PON2-KD cells (**Figure 5H-I**). Of note, in shPON2 PON2-KD cells, MPP^+^ treatment specifically increased cellular and mitochondrial ROS and reduced overall mitochondrial oxygen consumption profile, suggesting that PON2-KD cells may be more susceptible to MPP^+^-induced mitochondrial damage. Overall, these data show that vutiglabridin has significant neuroprotective effects against mitochondrial toxin and that PON2 is required for these effects.

**Fig 5.**
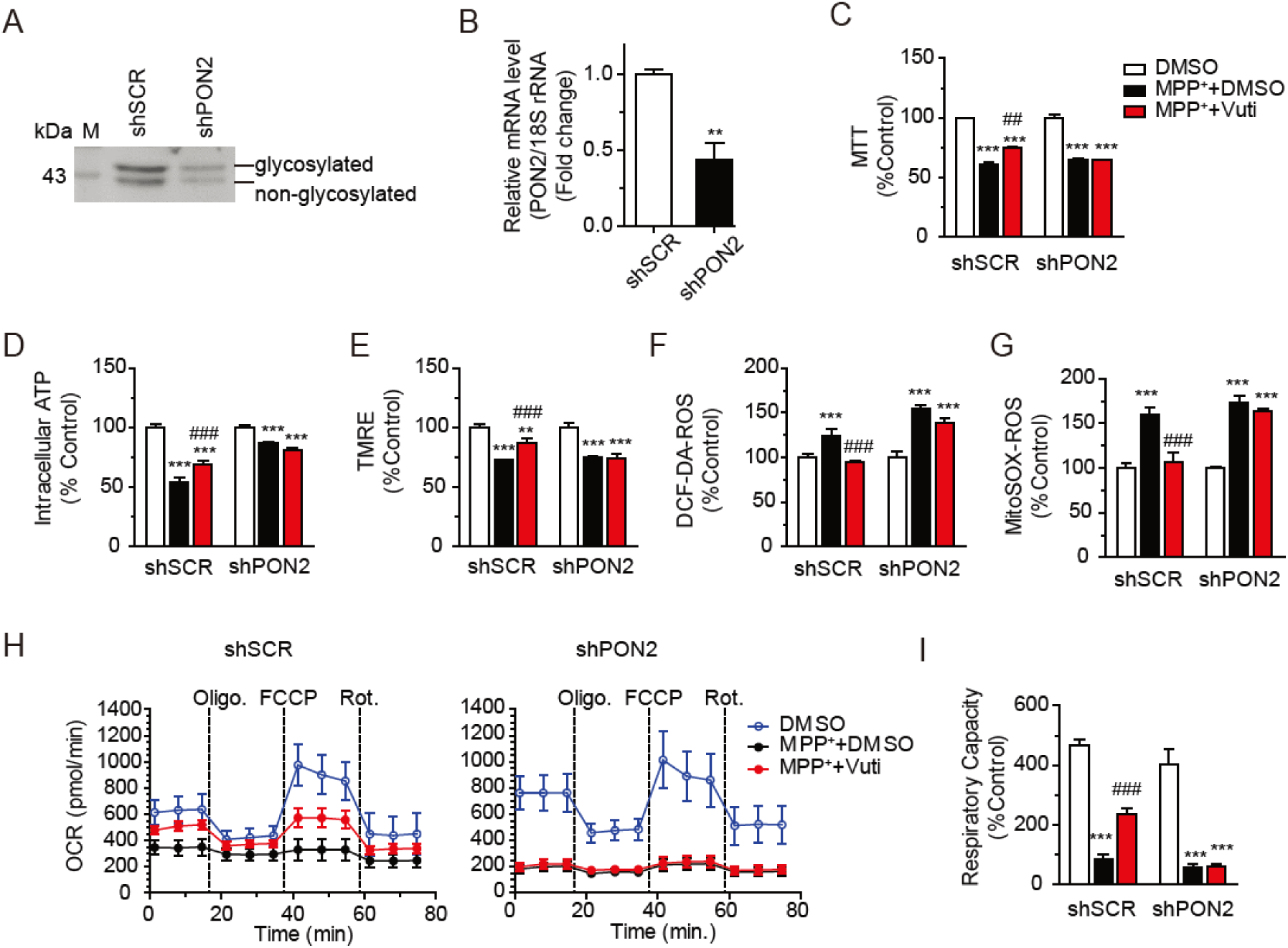
Vutiglabridin restores MPP^+^-induced damage in neuronal cells via PON2. PON2 knockdown (KD) SH-SY5Y cells were established by infecting with lenti-shPON2 viral particles. Verification of PON2 KD by (A) Western blot (B) real-time PCR. The values are reported in comparison to the control (shSCR). \*\**p* < 0.01 vs. Con, from a Student’s t-test. (C-G) shPON2-SH-SY5Y (shPON2) cells were placed in a 96-well plate and incubated with 1 mM MPP^+^ for 20 h, and then treated with 1 nM vutiglabridin (Vuti) for 24 h, after which the mitochondrial activity was analyzed. (C) MTT as NADH dehydrogenase activity of OXPHOS complex 1. (D) Intracellular ATP content. (E) TMRE-mediated mitochondrial membrane potential. (F) DCF-DA-based total ROS generation. (G) MitoSOX-based mitochondrial superoxide generation. All values are reported as a percentage of the Control. The data are plotted as the mean ± SEM (n = 6). (H, I) OCR. Cells in XF-24 microplate were incubated with 1 mM MPP^+^ for 20h, and then were treated with 1 nM vutiglabridin for 24 h. OCR was analyzed using a Seahorse XF-24 analyzer. Oligomycin, FCCP, and rotenone were consecutively injected to obtain mitochondrial respiratory capacities. (H) OCR profiles of shSCR control cells and shPON2 cells. (I) Total respiratory capacity (FCCP-induced OCR). The data are plotted as the mean ± SEM (n = 3). \**p* < 0.05, \*\**p* < 0.01, \*\*\**p* < 0.001 vs. Control (DMSO, white bar); ^#^*p* < 0.05, ^##^*p* < 0.01, ^###^*p* < 0.001 vs. MPP^+^ + DMSO (black bar). *p* values are from a one-way ANOVA followed by Tukey’s test.

### Knockdown of PON2 abolishes vutiglabridin-mediated effects on MPTP-injected mice

We investigated whether PON2 is also required for the in vivo neuroprotective effects of vutiglabridin. We performed the same set of animal study with the wildtype mice (WT) with an addition of PON2 knockdown (PON2 KD) mice group, which was generated by gene trapping in intron 2 and is reported to have a slight leakiness (<10%) of PON2 in the whole body (*19*). We found that in the brain of the PON2-KD mice, PON2 mRNA level is almost completely eliminated (**Figure 6A**). While 50 mg/kg of vutiglabridin treatment for three weeks was able to significantly protect against MPTP-induced damage to motor behaviors and dopaminergic neurons in wildtype mice, such neuroprotective effects were not observed in PON2-KD mice (**Figure 6C-H**). An additional group of 50 mg/kg of vutiglabridin treatment to non-MPTP-injected wildtype mice did not show any altered motor behavior or dopaminergic neuronal cell count and density compared to the vehicle-only treated group, which suggest that vutiglabridin itself does not induce any behavioral or neuronal change to wildtype. The efficacy observed in MPTP-injected mice is solely due to the effects of vutiglabridin against MPTP treatment. Overall, these data show that PON2 is required for the neuroprotective effects of vutiglabridin in MPTP-injected PD mice and that PON2 is indeed a key target protein of vutiglabridin.

**Fig 6.**
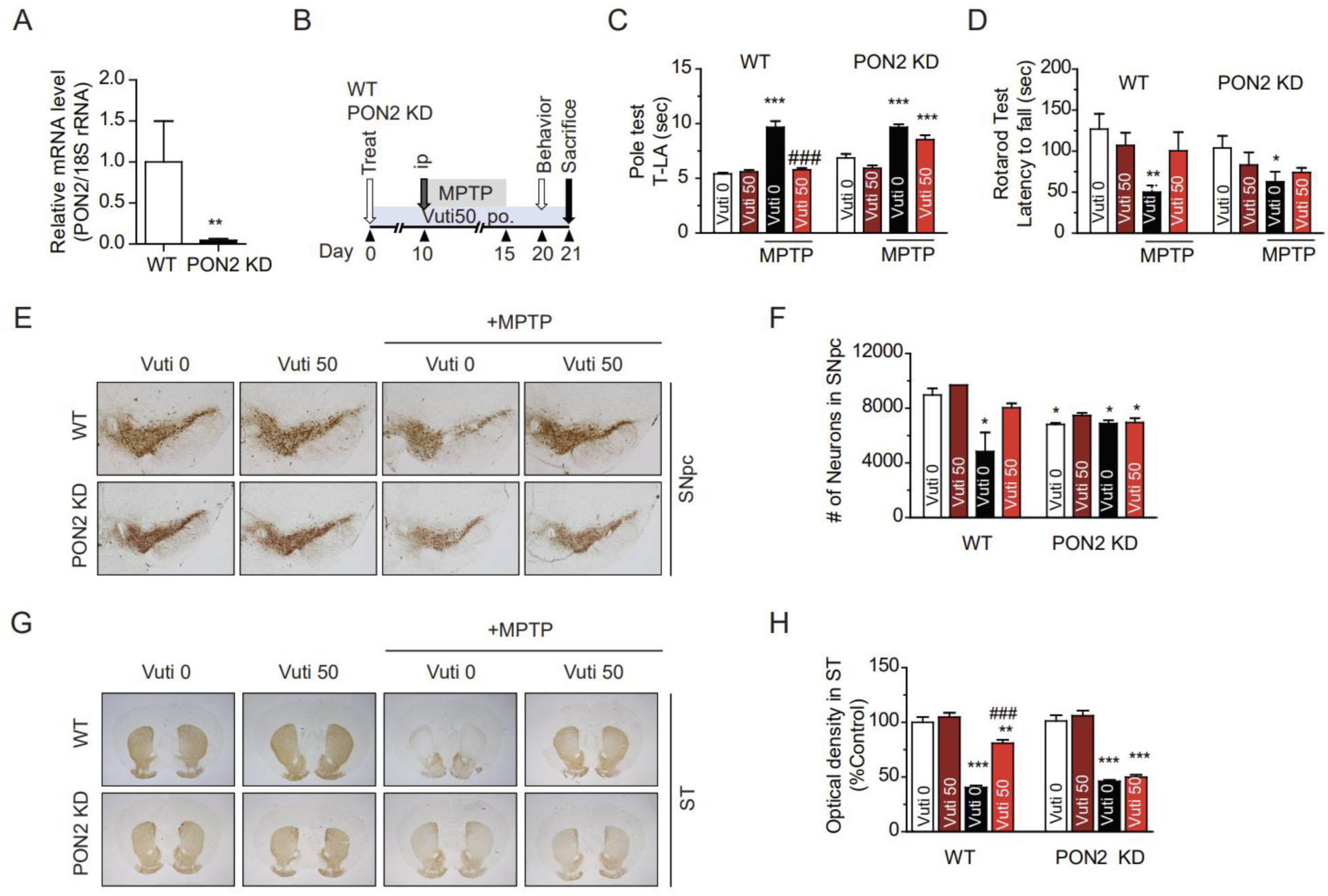
Vutiglabridin protects MPP^+^-induced damage in mice via PON2. (A) Realtime RT-PCR of PON2 mRNA level in the total brain homogenates of WT and PON2 KD mice. The data are plotted as the mean ± SEM (n = 4). \*\**p* < 0.01 vs. WT, from a Student’s t-Test. (B) Scheme of the experimental design. Male wildtype C57BL/6 mice (WT) and PON2-knockdown mice (KD) of 8-10 weeks of age were orally administered with 0 or 50 mg/kg of vutiglabridin (Vuti 0 and Vuti 50) for 21 consecutive days (n=5-8). All groups except for the vehicle-only treated group were injected with MPTP intraperitoneally at 30 mg/kg for five days starting on day 10. All compounds were administered daily. On day 20, the rotarod and pole test were performed. On day 21, mice were sacrificed, and dopaminergic neurons were visualized via tyrosine hydroxylase (TH) immunohistochemistry. (C) Latency time (sec) to arrive at the floor (T-LA) in the pole test. (D) Latency time (sec) in the rotarod test. (E) Representative photomicrographs of the SNpc. (F) Stereological count of the number of TH-immunopositive neurons in SNpc. (G) Representative photomicrographs of the striatum (ST). (H) Optical density in the ST. The data are plotted as the mean ± SEM (n = 5-8). The number of animals for each group is specified in the methods. \**p* < 0.05, \*\*\**p* < 0.001 vs. Vehicle group (Vuti 0 without MPTP); ^#^*p* < 0.05, ^###^*p* < 0.001 vs. MPTP + Vehicle group (Vuti 0). *p* values are from a one-way ANOVA followed by Tukey’s test.

### Assessment of neurotoxicity by in vitro safety screening and FOB evaluation

The overall safety of vutiglabridin has been established in human phase 1 clinical trials (NCT04732988, NCT04733001, NCT04703764) (*45*). However, to be safely repositioned as a central nervous system (CNS)-targeting drug candidate, potential neurotoxicity of vutiglabridin was assessed through a comprehensive in vitro safety screening. Vutiglabridin interaction was tested to an extensive total of 225 mammalian proteins including ion channels, G-protein coupled receptors (GPCR), nuclear receptors, kinases, non-kinase enzymes, and drug transporters (**Supplemental Table S2 and S3**). In addition to assessing neurotoxicity, these panels also included assessment of cardiac function and drug-drug interaction – since most PD patients are over 60 years old and may have other health complications – and assessment of the reported direct targets of glabridin (*26*) – estrogen receptor, peroxisome proliferator-activated receptor gamma (PPARγ), and cyclooxygenase 2 (COX-2) – to examine whether vutiglabridin interacts with those target proteins. We found that vutiglabridin does not interact with all of the assayed proteins, under the cutoff value of IC_50_/EC_50_ of 10 μM, except for the following three: cholecystokinin CCK1 receptor (CCK_A_; IC_50_ = 3.84 μM), melatonin receptor MT2 (IC_50_ = 3.78 μM), and breast cancer resistance protein (BCRP; IC_50_ = 8.18 μM). These findings show that vutiglabridin does not bind to the reported targets of glabridin other than PON2 and also demonstrate robust in vitro safety of vutiglabridin.

To further investigate potential neurotoxicity of vutiglabridin in vivo, a standard functional observational battery (FOB) evaluation (*46*) was performed. Male Sprague-Dawley were administered once with very high doses of vutiglabridin – 500, 1000, and 2000 mg/kg – and two observers blindly scored their observations of the animals in home cage, open field, hand-held, and sensory motor function at 0.5, 1, 3, 6, and 24 h timepoints. There were no statistically significant changes induced by vutiglabridin in all observations (**Supplemental Table S4**), demonstrating the preclinical safety of vutiglabridin in the context of neurotoxicity.

## DISCUSSION

Clinical development of safe and efficacious pharmacotherapies that slow the progression of PD is strongly needed, while novel target proteins that ameliorate mitochondrial dysfunction provide promising directions for the development of such treatments for PD. In this study, we show that vutiglabridin as a clinical-stage compound modulates PON2 to induce clear neuroprotective effects in preclinical models of PD. Vutiglabridin was also found to be safe in a comprehensive safety screening on CNS. We make a strong case for PON2 as a potential novel target protein for the development of treatments for PD and for the clinical development of vutiglabridin.

We first show that vutiglabridin crosses the BBB. Quantitative autoradiography has been historically employed to evaluate drug transport to the brain (*47*); while the strength of the technique lies in the high spatial resolution to potentially evaluate which areas in the brain the radioactive compound is located (*48*), the limit of our experiment was that whole-body sagittal sections were made instead of brain-specific coronal section, so there was low resolution. Still, the radioactivity of vutiglabridin in rat was clearly observed throughout all regions of the brain at a level 2.5-fold compared to the radioactivity in the blood, and this value was remarkably replicated in the pharmacokinetics study in blood and brain of mice, which consistently and strongly show that vutiglabridin penetrates BBB. Limited brain penetration has been a key hurdle to the development of many CNS-targeted pharmacotherapies. Glabridin has also been reported to have limited brain penetration as it is a substrate of P-gp (*34*), yet even with this limited permeability, glabridin improved memory function in mice (*28*) and in diabetic rats (*29*). Vutiglabridin had structural modifications that likely have contributed to its enhanced brain penetration, as it no longer is a P-gp substrate, and therefore holds more promise as a neuroprotective agent.

PON2 is a protein that needs to be studied more in depth. In this study, we created PON2 3D structure via homology modeling using PON1 and purifiable full-length recombinant PON2 with His-Tag at the C-terminal, both of which may aid future research on PON2. Using the in silico model, we found that both (R) and (S)-form of vutiglabridin binds to PON2; this is an exceptional instance where both enantiomers of a compound bind to the same target protein and thereby supports the development of vutiglabridin as a racemate. Also, compared to glabridin, vutiglabridin had greater binding interactions; the hydrogenation of double bond in pyranobenzene and the ethoxification in the resorcinol ring of vutiglabridin allow greater conformational flexibility and hydrophobicity (*25*), which may explain its increased alignment in both hydrogen bonding (ILE57, VAL335) and hydrophobic interaction (PRO59, TYR336) with PON2. While we found that vutiglabridin binding to PON2 increases its protein stability, the exact molecular mechanism of such effect requires more investigation.

We found that vutiglabridin protects against MPP^+^-induced mitochondrial dysfunction in neuronal cells via PON2. The exact mechanism of PON2 in reducing mitochondrial ROS is yet unknown; while there is speculation that it binds to coenzyme Q_10_ at its semiquinone form to prevent superoxide generation (*22*), such direct molecular function in relation to the molecular structure of PON2 has not been elucidated. Of note, vutiglabridin was able to restore MPP^+^-induced mitochondrial damage as a post-damage treatment; therefore, it is possible that vutiglabridin may function to induce mitophagy for the clearance of already-damaged mitochondria. PON2 was previously reported to interact with protein deglycase DJ-1, also known as Parkinson disease protein 7 (PARK-7), whose mutations are associated with early-onset PD (*49*). DJ-1 promoted PON2 activity to enhance neuronal survival against oxidative stress in cells (*50*), and DJ-1 is reported to act in parallel with PINK1/parkin pathway to induce mitophagy (*51*). Future research may investigate the interaction between PON2, DJ-1, and vutiglabridin in affecting mitophagy and mitochondrial dynamics.

Vutiglabridin showed neuroprotective effects in MPTP-injected subacute model of PD, and such effects were abrogated in PON2-KD mice. A limitation within our studies was the significant number of deaths, to about 50% of the group, induced by MPTP injection in PON2-KD mice compared to only 20% in wildtype mice, which suggest that PON2-KD mice may be more susceptible to the neurotoxin. In PON2-KD mice, the number of TH-positive neurons in SNpc was not significantly reduced (**Figure 6E, F**), while the motor behavior function and the optical density of TH in striatum were significantly reduced by MPTP injection. One can surmise that only PON2-KD mice that received less damage to SNpc survived to be analyzed at sacrifice, although no evidence to support such claim could be acquired. Regardless of such limitation, there was clear abrogation of the neuroprotective effects of vutiglabridin in PON2-KD mice, and vutiglabridin treatment also did not change the survival rate of MPTP-injected PON2-KD mice. Our findings show that vutiglabridin acts on PON2 to protect mice from MPTP-induced neuronal damage. We report for the first time a compound that modulates PON2 to provide neuroprotection against a mitochondrial toxin in PD model.

Because vutiglabridin was developed based on a phenotypic screening approach, as opposed to a target-based approach, an examination of its potential neurotoxicity by an extensive study of its potential off-targets was warranted. This study provides such information and demonstrates strong CNS-related safety. Also, an examination of potential targets based on the known targets of glabridin was performed; while the structural modifications made to vutiglabridin enhanced its binding to PON2, they have also dismissed other proteins as targets – estrogen receptor, PPARγ, and COX-2 – which may facilitate the mechanistic understanding of vutiglabridin as a less promiscuous compound. Based on the findings of the in vitro safety panels, clinical evaluation of potential interaction effects on cholecystokinin CCK1 receptor, whose signal acts as a hunger suppressant (*52*), and melatonin MT2 receptor, which modulates sleep and circadian phase (*53*), should occur by monitoring changes in patient food intake and sleep schedule, both of which are crucial aspects in PD patients. Also, co-administration with breast cancer resistance protein (BCRP)-mediating drugs such as pitavastatin and rosuvastatin (*54*) should be heeded during patient recruitment. The absence of any findings in the FOB evaluation at doses 40-fold higher than the maximum efficacy dose alleviate concerns on neurotoxicity.

In conclusion, this study provides preclinical basis for the clinical development of vutiglabridin and proposes the modulation of PON2 as a therapeutic approach for treating PD. The findings in this study, along with the clinical data acquired from the ongoing phase 2 study in obesity, will support immediate proof-of-concept study of vutiglabridin in PD patients, and if successful, can provide a new gateway of drugs targeting PON2 for ameliorating mitochondrial dysfunction in PD.

## MATERIALS AND METHODS

### Study Design

The goal of the study was to evaluate potential neuroprotective effects of vutiglabridin in toxin models of PD and investigate whether such effects are mediated by PON2. SH-SY5Y human neuroblastoma cell line, Sprague-Dawley rats, and C57BL/6J mice were used in this study. QWBA study was performed in SD rat in accordance with established standard procedure (*38*) where we measured plasma pharmacokinetics first, and then acquired sections from one rat at each of the timepoint selected based on the plasma pharmacokinetics. In vitro assays were performed with three independent replicates, except for the comprehensive safety panels, where only two replicates were performed considering the nature of screening purpose. For animal studies of efficacy, we have not calculated sample size via power analysis, but we aimed to have greater than five animals per group, except for the pharmacokinetics study where we deemed three animals per group to suffice because animals were exorbitantly sacrificed as per timepoint basis. For the efficacy study, three-week drug administration and five-day MPTP injection have been previously optimized, as well as the behavioral and biological endpoints (*55–57*). There were unexpected deaths of PON2-KD animals during the five days of MPTP injection, and they were excluded from analysis. Details on randomization, blinding, animal age and sex, and example sample size are written under each relevant figure and/or methods.

### Materials

Vutiglabridin was prepared at Glaceum Inc. (Suwon, Korea) following the protocols from Patent US9783551B2 (*58*). Primary antibodies against TH (Cell Signaling Technology, Beverly, MA, USA) and β-actin (Sigma-Aldrich Co., St. Louis, MO, USA) were purchased from commercial sources. Tetramethylrhodamineethylester (TMRE), 5,6-chloromethyl-2’,7’-dichlorodihydrofluorescein diacetate acetyl ester (DCF-DA), and MitoSOX agents were purchased from Molecular Probes (Eugene, OR, USA). All other reagents were purchased from Sigma-Aldrich. Media and culture reagents were products of Gibco Industries Inc. (Auckland, New Zealand).

### Quantitative whole-body autoradiography (QWBA) assay in rat

Male Sprague Dawley rats (n = 5, 16 h fasted; Charles River Laboratories Japan Inc., Yokohama, Japan) of 8 weeks of age were administered 10 mg/kg of ^14^C-labeled vutiglabridin (Glaceum Inc., Suwon, Korea) – equivalent to radioactivity of 3.7 MBq/kg – through oral gavage. After 2, 6, and 24 h, each rat was euthanized by CO_2_ inhalation. Its nasal cavity and anus were filled with 4% w/v CMC-Na, and the carcass was frozen in a dry ice-acetone mixture. A 30-µm thick whole-body sagittal sections were cut with a cryomicrotome (CM3600, Leica Biosystems, Wetzlar, Germany), freeze-dried, covered with 4-µm thick protective membrane (Diafoil, Mitsubishi Plastics, Tokyo, Japan), and then exposed to the imaging plate (TYPE BAS SR2040, Fujifilm, Tokyo, Japan) for 24 h in a sealed lead box along with the plastic standard samples (CFQ7601, Amersham Biosciences Corp., NJ, USA) for calibration. The radioactivity was measured by the bio-imaging analyzer system (FUJIX-BAS2500, Fujifilm, Tokyo, Japan; resolution µm, gradation 256, sensitivity 10000, latitude 4) where the radioactivity in organs was converted to the photo-stimulated luminescence per unit area (PSL/mm^2^) minus the background radioactivity of the plastic standard samples. Tissue radioactivity in the whole-body autoradiograms were quantified by densitometry using an MCID image analysis software (v. 7.0, MCID Image Analysis Software Solutions for Life Sciences, Cambridge, UK). Radioactivity concentrations were expressed as ng equivalents of ^14^C-vutiglabridin per gram tissue (ng eq/g tissue). The study protocol references numerous established methods for QWBA (*38, 47, 59*) and was reviewed and approved by the Institutional Animal Care and Use Committee (IACUC: 2017-016).

### LC-MS/MS analysis

LC-MS/MS analysis to quantify vutiglabridin used previously set conditions (*60*). Briefly, HPLC-ESI-MS/MS analyses were performed using a 1200 series HPLC system (Agilent Technologies, Wilmington, DE) coupled to 6490 Accurate-Mass Triple Quadrupole Mass Spectrometer (Agilent Technologies, Wilmington, DE). Mass detection was performed in the positive ion mode, and the column temperature was maintained at 40°C using a thermostatically controlled column oven. The column used for the separation was Hypersil GOLD C18 column (2.1 × 100 mm ID; 1.9 μm, Thermo Science). The mobile phases consisted of 0.1% acetic acid in D.W (solvent A) and 0.1% acetic acid in ACN:MeOH=3:1 (v/v) (solvent B). The LC gradient was as follows: 0−3 min, B 20%; 3−5 min, B 20−80%; 5−8 min, B 80%; 8−10 min, B 80−20%; and 10−15 min, B 20%. The flow rate was 250 μL/ min, and total run time was 15 min. In multiple reaction monitoring (MRM) analyses, the target transition used were 355.1/151.0 m/z for vutiglabridin. The collision energy was 17 V and nitrogen was used as the desolvation gas at a flow rate of 11 L/min and at 300°C. Skyline software (MacCoss Laboratory, University of Washington, Seattle, WA) was used to process the LC-MS/MS data.

### Quantification of drug concentration in the plasma and brain of mice

8-week-old male C57BL/6J mice (n = 24, non-fasted; Charles River Japan (CRJ), OrientBio, Seongnam, Korea) were dosed with 50 mg/kg of vutiglabridin once and sacrificed at 0, 1, 2, 4, 6, 8, 10, 24 h timepoint (n = 3 per timepoint). Their whole blood was extracted via vena cava in herapin-coated syringe and their whole brain was extracted after perfusion with 0.9% saline. Vutiglabridin was quantified in both plasma and brain samples using LC-MS/MS analysis. Study was performed in accordance with the Association for Assessment and Accreditation of Laboratory Animal Care International (AAALAC; Approval No. LCDI-2021-0027).

### PON2 3D modeling and molecular docking study

The 3D structure of PON2 was created by prediction using homology modelling of its family protein PON1, whose 3D structure is known and there is sequence identity of 61.7% and sequence similarity of 79.2%. The structure of PON1 (PDB ID: 1V04) was obtained from the RCSB Protein Data Bank (http://www.rcsb.org). The 3D structure of PON2 was predicted using Discovery Studio (DS) 2018 (BIOVIA, San Diego, CA, USA). The structural stability of the constructed model of the PON2 protein was verified through molecular dynamic simulation, which was run for a total of 10 ns, and RMSD and RMSF were investigated to determine system stability. Phi (Φ) and psi (Ψ) torsion angles and interatomic collisions were reviewed for all amino acids through the Ramachandran plot. Molecular docking calculation was carried out by Genetic Optimization for Ligand Docking (GOLD v5.2.2, The Cambridge Crystallographic Data Centre, Cambridge, UK), an automated docking program to predict the binding mode of the ligand by genetic algorithm (*61, 62*). The geometry of the compounds – (R) and (S) form of vutiglabridin and glabridin – was optimized by energy minimization through Minimize Ligands tool in DS. All the active site residues within 10 Å radius sphere of the center were included for the calculation. The number of docking runs was set to 50 for each compound. All other parameters were set as default.

### Mass spectrometry-based binding assay

The method followed the concept of previously established protocol (*41*). Briefly, recombinant rePON2 was diluted with buffer (Tris–HCl pH 7.6 50 mM, CaCl_2_, 10 mM), and vutiglabridin (10 nM) was co-incubated with varying concentrations of PON2 (0, 5, 10, 20, 50, 100, 200, and 500 nM) for 30 min at 30 °C. PON2 was separated using 7K size-exclusion chromatography column (Thermo Fisher Scientific) and was incubated with a denaturing buffer (1% formic acid in acetonitrile) at 80°C for 10 min. The supernatant, which now contains PON2-bound vutiglabridin, was collected, dried, and re-suspended in acetonitrile, and the drug quantification was performed via LC-MS/MS analysis.

### Thermal shift assay

Recombinant rePON2 (500 nM) and 10 nM vutiglabridin were incubated for 30 min in tube under the following temperatures: 37, 42, 47, 52, 57, 62, 67 ⁰C. The supernatant after centrifugation was collected as non-degraded PON2, and its quantity was measured via LC-MS analysis.

### Cell culture and treatment

Human SH-SY5Y neuroblastoma cells (ATCC® (CRL-2266™; Manassas, VA, USA) were first cultured in Dulbecco’s Modified Eagle Medium (DMEM)/F12 supplemented with 10% fetal bovine serum (FBS), 100 U/mL penicillin, and 100 μg/mL streptomycin (complete media, CM) at 37°C with 5% CO_2._ Cells (5×10^4^ cells/well) and then were cultured in 96-well plates for 24 h followed by incubation in serum-deficient media (SDM, culture media containing 0.5% FBS) for 16 h. Quiescent cells in SDM were incubated with 1 mM MPP^+^ or dimethyl sulfoxide (DMSO) vehicle for 24 h, then treated with vutiglabridin at the designated doses for 24 h. The treated cells were harvested according to the purpose and analyzed.

### PON2 knockdown using shPON2

Lentiviral PON2 shRNA (h) (sc-62838) was purchased from Santa Cruz Co. (Santa Cruz, CA, USA). shRNA lentiviral particles transduction was performed as manufacturer’s protocol. Total RNA was isolated with Trizol reagent (Invitrogen, Carlsbad, CA, USA). Total RNA (1.5 μg) was reverse transcribed using MMLV reverse transcriptase (Promega, Madison, WI, USA), RNasin Ribonuclease inhibitors (Promega), 10 ng of random primers (Invitrogen), and 25 mM of dNTP mix (Gene Craft, Ludinghausen, Germany), according to the manufacturer’s instructions. Real-time quantitative RT-PCR (qRT-PCR) was performed using primers for human PON2 (5’-CCA AGC AAG GGA CAG AAA AG −3’ and 5’-TCA CAG TGC CAG AAG TGA GG −3’) and 18S rRNA (5’-GAG CGA AAG CAT TTG CCA AG-3’ and 5’-GGC ATC GTT TAT GGT CGG AA-3’) on a Roter-Gene Q (Qiagen, Hilden, Germany) with 2x AmpiGene® qPCR Green Mix Lo-ROX (Enzo Biochem, NY) at 95°C for 10 min, followed by 45 cycles of 95°C for 5 s and 60°C for 15 s and 72°C for 20 sec. Measurements were performed in duplicate for each sample. The quantity of mRNA was corrected by simultaneous measurement of nuclear DNA encoding 18S rRNA. The relative quantification in gene expression was determined using the 2-ΔΔCt method. Relative levels of mRNA expression are presented as fold changes compared to those under the control condition.

### Cell-based mitochondrial activity assays

A systemic cell-based mitochondrial function analysis system was established using SH-SY5Y cells in 96-well plates based on fluorescence detection methods as described previously (*9, 57*). The assay system covers quantitative assays for methyl thiazyl tetrazolium-mitochondrial dehydrogenase activity (MTT), TMRE-based mitochondrial membrane potential, ATP contents, and ROS of cells. Cells in black 96-well culture plate were incubated with 200 nM TMRE and 0.5 μM Hoechst 33342 for 30 min at 37°C in phenol red-free SDM, or with 1 μM CM-H_2_DCFDA or 5 μM MitoSOX and 0.5 μM Hoechst 33342 for 1 h at 37°C. Fluorescence intensities at 550 nm/580 nm for TMRE, at 510 nm/580 nm MitoSOX, or at 494 nm/522 nm for DCF-DA were normalized by Hoechst intensity at 355 nm/480 nm (Spectramax Gemini EM, Molecular Devices, Sunnyvale, CA, USA).

Endogenous OCR was measured in adherent SH-SY5Y cells using a Seahorse XF-24 Analyzer (Seahorse Bioscience, Billerica, MA, USA) following the manufacturer’s protocol with minor modifications (*63*). Briefly, OCR assays were performed using SH-SY5Y cells (5.0 × 10^3^ cells/well) equilibrated in DMEM without sodium bicarbonate in XF-24 microplates. After measuring basal OCR for 3 min, oligomycin (1 μg/mL), carbonyl cyanide-p-trifluoromethoxyphenylhydrazone (FCCP) (0.3 μM), and rotenone (0.1 μM) were consecutively added into each well to reach their final working concentrations. The OCR was calculated from 3-min measurement cycles and normalized to cell number.

### Western blot analysis

Cell lysates were prepared on ice in PRO-PREP lysis buffer (10 mM HEPES, pH 7.9, 10 mM KCl, 2 mM MgCl_2_, 0.5 mM dithiothreitol, 1 mM phenylmethylsulfonyl fluoride, 1 μg/mL aprotinin, 1 μg/mL pepstatin A, and 2 μg/mL leupeptin; iNtRON Biotechnology, Gyeonggi-do, Korea). Protein extracts (30 μg) were separated by 12% SDS-PAGE and analyzed by Western blot and an enhanced chemiluminescence system (ECL, Amersham Bioscience, Piscataway, NJ, USA). Anti-PON2 antibody (Santa Cruz Biotech, Dallas, TX, USA) and anti-TH (Pel-Freez Biologicals, Rogers, AR, USA) antibodies were diluted at 1:1000. Anti-β-actin antibody (1:3000) was used as a loading control. Band intensities were quantified by densitometry and ImageJ program (National Institutes of Health, Bethesda, MD, USA).

### Quantitative real-time PCR analysis

PON2 level was quantified in the mouse whole brain homogenate in 100 mg per 1 ml of QIAzol® lysis reagent (Qiagen, Hilden, Germany). mRNA 500 ng was used for cDNA synthesis in 10 μl volume according to the manufacturer’s instructions of PrimeScript^TM^ RT-PCR kit (Takara Bio Inc., Shiga, Japan). Real-time RT-PCR was then performed using the KAPA SYBR FAST qPCR Master Mix (2x) Kit (Kapa Biosystems, Cape Town, South Africa) in LightCycler 480® (Roche, Basel, Switzerland) on 384 well with the final cDNA concentration of 10 ng in 10 μl reaction volume. The primers used were mouse PON2 (F-ACCCACTTCTACGCCACCA, R-CTGCCACCAGTTTAACTTCTTCT) obtained from Primer bank (ID: 34303979a1) and mouse Rn18s (F-AGTCCCTGCCCTTTGTACACA, R-CGTTCCGAGGGCCTCACT). The PCR conditions were 95 °C for 3 min, 40 cycles of 95 °C for 3 s and 60 °C for 30 s, and 8 °C hold. Data was analyzed by converting 2^−ΔΔ*Ct*^, normalized by Rn18s mRNA expression.

### MPTP-injected mice and drug administration

Male C57BL/6 mice aged 8–10 weeks with initial body weight of 20-24 g were purchased from Samtako Bio Korea Co. Ltd (Osan, Korea) and PON2-KD mice were generously provided by Dr. Srinuvasa T. Reddy (UCLA, CA, USA). Animals were allowed at least 1 week of acclimatization before being subjected to the study and were housed in a regulated environment with a 12 h/12 h light/dark cycle. Food and water were provided *ad libitum*. The animal experiment was carried out in accordance with the National Institutes of Health Guide for the Care and Use of Laboratory Animals (NIH Publications No. 80–23; revised 1996). All procedures for handling mice were carried out in accordance with “Principles of Laboratory Animal Care” and the Animal Care and Use guidelines of Kyung Hee University, Seoul, Korea (KHSASP-20-163).

MPTP was dissolved in distilled water and injected intraperitoneally for five days from day 10 of the 21-day experiment period. Vutiglabridin or rasagiline were administered via oral gavage for 21 days. Vehicles for both vutiglabridin and MPTP (distilled water) were administered to all mice. Mice were randomly divided for all studies. For the first set of study shown in Figure 3, each group consisted of 5 animals. For the second set of study shown in Figure 6, each group consisted of the following animals: (1) wildtype vehicle-only group (Vuti 0; n = 5), (2) wildtype vutiglabridin 50 mg/kg group (Vuti 50; n = 5), (3) wildtype MPTP + vehicle-only group (Vuti 0; n = 8, excluding 2 deaths), (4) wildtype MPTP + vutiglabridin 50 mg/kg group (Vuti 50; n = 6), (5) PON2-KD vehicle-only group (Vuti 0; n = 7), (6) PON2-KD vutiglabridin 50 mg/kg group (Vuti 50; n = 4), (7) PON2-KD MPTP + vehicle group (Vuti 0; n = 13, 6 deaths excluded from analysis), (8) PON2-KD MPTP + vutiglabridin 50 mg/kg group (Vuti 50; n = 14, 5 deaths excluded from analysis).

### Animal behavior test

To determine forelimb and hindlimb motor coordination and balance, we performed the rotarod test as described previously (*64, 65*). Also, the pole test was performed to measure bradykinesia. For the rotarod test, the animals were pre-trained on the rotating bar of a rotarod unit set (LE 8500, Letica, Spain) on days 18 and 19 before the test on day 20. During pre-training, three trials per day were performed (5 rpm rotation speed on the first day, 15 rpm rotation speed on the second day). Mice were kept on the rotating bar (7.3-cm diameter) for 5 min on each trial. On day 20, the time spent on the rotating bar at 20 rpm, which was defined as the latent period, was recorded. Performance was recorded as 300 s if the latent period exceeded 300 s. Mice had at least 5 min of rest between trials to reduce stress and fatigue. Each animal underwent three test trials, and the mean of the test results was subjected to statistical analysis. For the pole test, mice were held on the top of the pole (diameter 8 mm, height 55 cm, with a rough surface). The time to land down and place four fee on the floor was recorded as the time for locomotion activity (T-LA).

### Immunohistochemistry of TH-positive neurons

On day 21, mice were anesthetized with sodium pentobarbital (50 mg/kg intraperitoneally) and transcardially perfused with a saline solution containing 0.5% sodium nitrate and heparin (10 U/mL) and then fixed with 4% paraformaldehyde dissolved in 0.1 M phosphate-buffered saline (PBS). Brains were dissected from the skull, post-fixed overnight in buffered 4% paraformaldehyde at 4°C and stored in a 30% sucrose solution until they sank. Subsequently, the brains were prepared for frozen section on Cryostat (Microsystems AG, Leica, Wetzlar, Germany) in 30-µm thick coronal sections, and rinsed in PBS. The tissue sections were incubated overnight at 4°C with primary rabbit anti-TH antibodies followed by staining with biotinylated anti-rabbit IgG and an avidin-biotin peroxidase complex (ABC) standard kit (Vector Laboratories, Burlingame, CA, USA). Signals were detected by incubating sections with 0.5 mg/ml 3,3′-diaminobenzidine (Sigma, St. Louis, MO, USA) in 0.1 M PBS containing 0.003% H_2_O_2_. After the labeled tissue sections were mounted on gelatin-coated slides, stained brain sections were imaged under a bright-field microscope (Olympus Optical, Tokyo, Japan). The TH-immunopositive cells in the SNpc in brain tissue sections (5 sections/series) were counted at x100 magnification. The TH-immunopositivity in ST was measured by the optical density of TH-positive fibers at x40 magnification using ImageJ software (National Institutes of Health, Bethesda, MD, USA).

### Stereological cell counting

The unbiased stereological estimation of the total number of TH-immunopositive neurons in the SN was made using the optical fractionator method performed on an Olympus computer-assisted stereological toolbox system version 2.1.4 (Olympus) as previously described (*66, 67*). The sections used for counting covered the entire SN from the rostral tip of the pars compacta (SNpc) back to the caudal end of the pars reticulata (SNr) (anterioposterior, −2.06 to −4.16 mm from the bregma). The SN was delineated on a x1.25 objective and generated counting grid of 150×150 μm. An unbiased counting frame of known area (47.87 x 36.19 μm = 1733 μm^2^) superimposed on the image was placed randomly on the first counting area and systemically moved through all the counting areas until the entire delineated area was sampled. The actual counting was performed using a x100 objective. The estimate of the total number of neurons was calculated according to the Optical Fractionator Equation.

### Statistical analysis

The results are presented as the mean ± standard error of the mean (SEM). Statistical significances between experimental groups were evaluated by one-way ANOVA with Tukey post-hoc testing analysis using GraphPad Prism (version 9.4.0, GraphPad Software, San Diego, CA, USA), unless otherwise noted. Values of *p* < 0.05 were considered statistically significant.

## Acknowledgments

We thank Dr. Srinuvasa T. Reddy for generously providing PON2-KD mice for this research. We acknowledge Sekisui Medical Co., Ltd for QWBA assays and the pharmacokinetics study in rats, Pharma Resources Shanghai for P-glycoprotein efflux assay, Eurofin Discovery for in vitro safety panel assays, Solvo Biotechnology for transporter assays, and Biotoxtech Co., Ltd for FOB examination.

## Author contributions

Conceptualization: LSC, HJJ, SY, YKP

Methodology: SL, KWL, JHP, JL, HJJ

Investigation: SK, SL, SI, JHK, DHK, MP, SKP, HML

Funding acquisition: SY, YKP

Project administration: SK, LSC, HJJ, HSP, YKP

Supervision: LSC, HSP, SY, YKP

Writing – original draft: SK, LSC, KWL, YKP

Writing – review & editing: LSC, YKP

## Data and materials availability

All data associated with this study are present in the main text or the supplementary materials. Information on the 3D structure of PON2 can be obtained by contacting Keun Woo Lee at. Vutiglabridin can be obtained through a material transfer agreement from Glaceum by contacting Leo Sungwong Choi at.

## Supplementary Materials

### Materials and Methods

#### Octanol-water partition coefficient measurement

1000 μg/mL of vutiglabridin was mixed with octanol, and an aliquot of 2 mL was mixed with the same amount of aliquot of 2 mL of pH7.4 phosphate buffer. The mixed solution was shaken at room temperature for 24 h, and the aqueous and octanol phase was separately analyzed for vutiglabridin concentration via HLPC. The limit of quantification (LOQ) was 0.2012 μg/ml. Three replicates were performed. Because no concentration was detected in the aqueous phase, LOQ value was used; the actual concentration in the aqueous phase is expected to be lower than LOQ. The octanol-water partition coefficient Log D was calculated as log of concentration in octanol phase divided by concentration in aqueous phase.

#### P-glycoprotein (P-gp) efflux assay

P-gp efflux ratio was determined in MDCKII-hMDR1, MDCKII (*68*), and Caco-2 cell (*69*) monolayer via LC-MS/MS analysis, in accordance with the standard FDA guideline on transporter-mediated drug interactions. 10 µM of GF120918 was used as P-gp inhibitor. The apical (A-to-B) and basolateral (B-to-A) permeability coefficient (P_app_) were measured to yield the efflux ratio.

#### Plasma concentration of ^14^C-labelled vutiglabridin

For measuring the radioactive concentration in the plasma, blood (250 uL) was collected from the tail vein of the animals (n = 3) of the exact same conditions as the animals used for QWBA assay. The plasma was separated by centrifugation (8000xg, 4 ℃, 5 min) into a scintillation vial, dissolved with 2mL of tissue solubilizer (Soluene-350, PerkinElmer Inc., MA, USA), and mixed with 10mL of scintillator (Hionic-Fluor, PerkinElmer Inc., MA, USA). The radioactivity (ng eg./mL) was measured using liquid scintillation counter (1900CA, 2500TR, 2700TR, PerkinElmer Inc., MA, USA) for 2 min, calibrated with the scintillator as the background sample. Then the tissue to blood ratio was calculated. The protocol was reviewed and approved by the Institutional Animal Care and Use Committee (IACUC: 2017-016).

#### Cloning, expression, and purification of recombinant *Hs*PON2

The codon-optimization of *Hs*PON2 was for *Escherichia coli* system and synthesized according to the full-length open reading frame (GenBank accession No. AAC41995.1) from National Center for Biotechnology Information (NCBI; http://www.ncbi.nlm.nih.gov). The *Hs*PON2 gene was amplified by a standard PCR method with a primer set (Forward; 5’-GCACTC<u>CATATG</u>GGGGCATGGGTCGGGTGTGGGTTGGCCGGGGATAGAGCCGGC TTTC-3’, Reverse; 5’-GCACTC<u>CTCGAG</u>AAGCTCGCAGTACAATGC-3’) and inserted into an expression vector (pET21b; New England Biolabs, USA) using *Nde*I and *Xho*I restriction enzyme sites (underlined). Notably, a six-His tag was fused to C-terminus of the protein to facilitate protein purification. The recombinant construct was transformed into *E. coli* BL21 (DE3) competent cells. The transformed cells were activated overnight and transferred to 1 L of LB broth (BD bioscience, USA) with 100 µg/mL ampicillin (Duchefa Biochemie, The Netherlands) containing 0.1% glucose, and then incubated at 37 ℃ and 200 rpm in incubator (Vision Scientific, Korea) until reaching the optical density of 0.6 - 0.8 at 600 nm (OD_600_).

The overexpression of the recombinant *Hs*PON2 protein was induced by the addition of 0.5 mM isopropyl β-D-1-thiogalactopyranoside (IPTG) (Duchefa Biochemie), and then further incubated for 7 h at 25℃ and 170 rpm. After the protein expression, the cells were harvested and resuspended in 30 mL of lysis buffer (10 mM Bicine, pH 9.0, 50 mM NaCl) and lysed for 4 min by sonication (30% Amplitude, 4 s sonication and 6 s rest) (Sonics and materials, USA). To obtain *Hs*PON2 forming inclusion bodies, the supernatant was removed by centrifugation (12,000 rpm, 4℃) for 20 min. The pellet was resuspended in 30 mL of UREA buffer (10 mM Bicine, pH 9.0, 50 mM NaCl, 4 M UREA). After that, the cellular debris was removed by centrifugation (12,000 rpm, 4℃) for 20 min. The supernatant was filtered by the 0.45 µm syringe filter (GVS, Italy). The protein refolding was conducted by the dialysis with buffer of 2 L (10 mM Bicine, pH 9.0, 50 mM NaCl, 1 mM CaCl_2_). The refolded *Hs*PON2 was purified using affinity chromatography with a 10 mL of Ni-NTA resin (QIAGEN, Germany) and an elution buffer (10 mM Bicine, pH 9.0, 50 mM NaCl, 250 mM Imidazole). The eluted *Hs*PON2 protein was analyzed by a 12% SDS-PAGE and concentrated up to approximately 1 mL by 10 kDa filter-size amicon (Merck Millipore, USA). For activity assay of *Hs*PON2, the sample buffer was changed into activity buffer (20 mM Bicine, pH 8.0, 1 mM CaCl_2_). Subsequently, the purified *Hs*PON2 was confirmed by using the western-blot assay with the anti-histidine antibody (MBL, England).

## Supplemental Figures and Tables

**Fig. S1.**
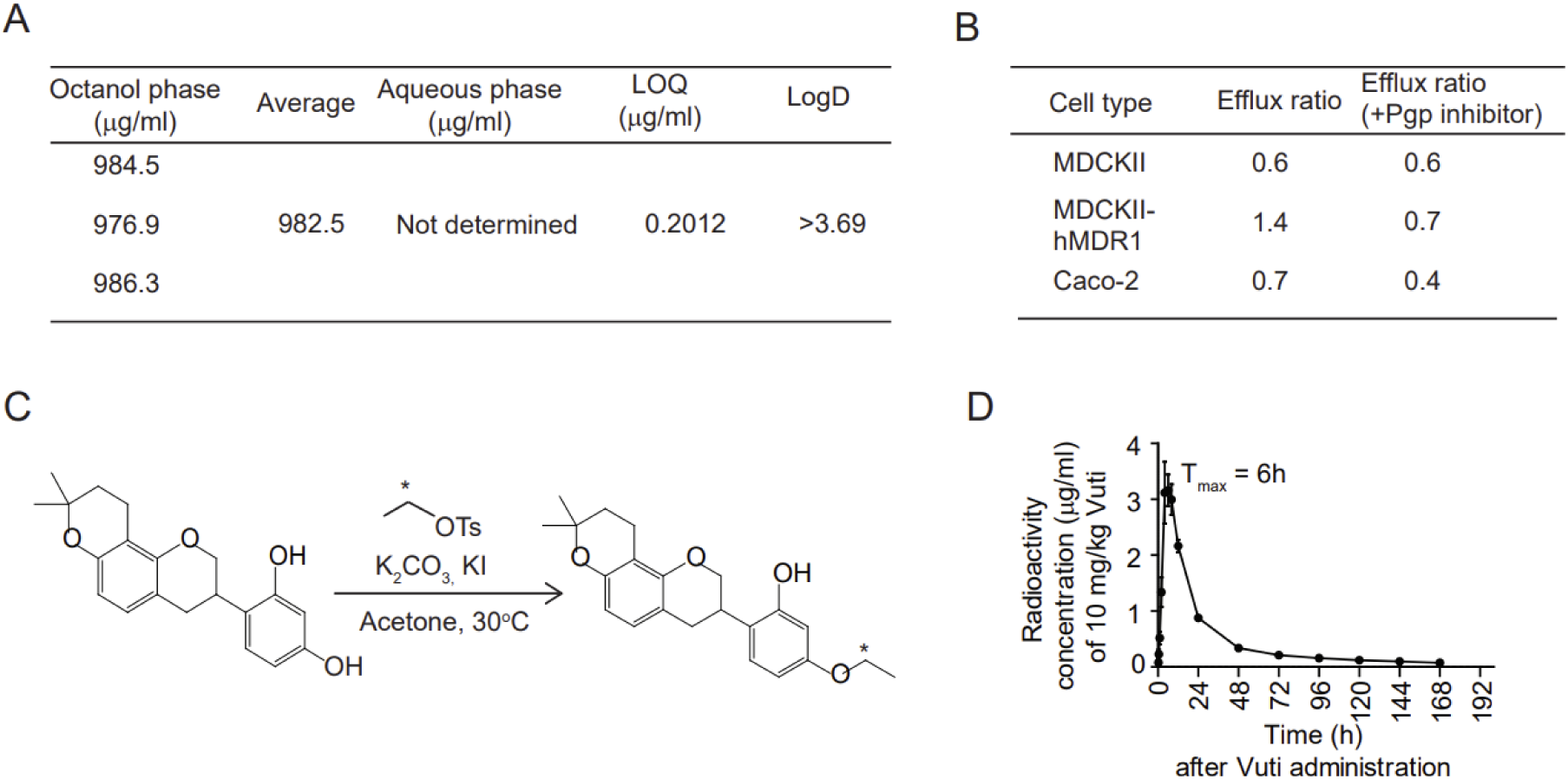
Properties of vutiglabridin and ^14^C-labeled vutiglabridin. (A) Measurement of vutiglabridin concentration in octanol and aqueous phase (pH7.4 phosphate buffer) via HPLC. Mean value of three replicates was used, and limit of quantification (LOQ) value was used to calculate Log D as log of concentration in octanol phase divided by concentration in aqueous phase. (B) Efflux ratios (P_app(B→A)_/P_app(A→B)_) of vutiglabridin in three types of cell monolayers, with or without P-glycoprotein (P-gp) inhibitor GF120918. (C) Schematic of ^14^C-labeling of vutiglabridin. Attachment to the ethoxy group is warranted because it is robustly maintained throughout metabolism. (D) Plasma radioactivity concentration in Sprague-Dawley rat after a single administration of 10 mg/kg of ^14^C-vutiglabridin (Vuti) (n = 3).

**Fig. S2.**
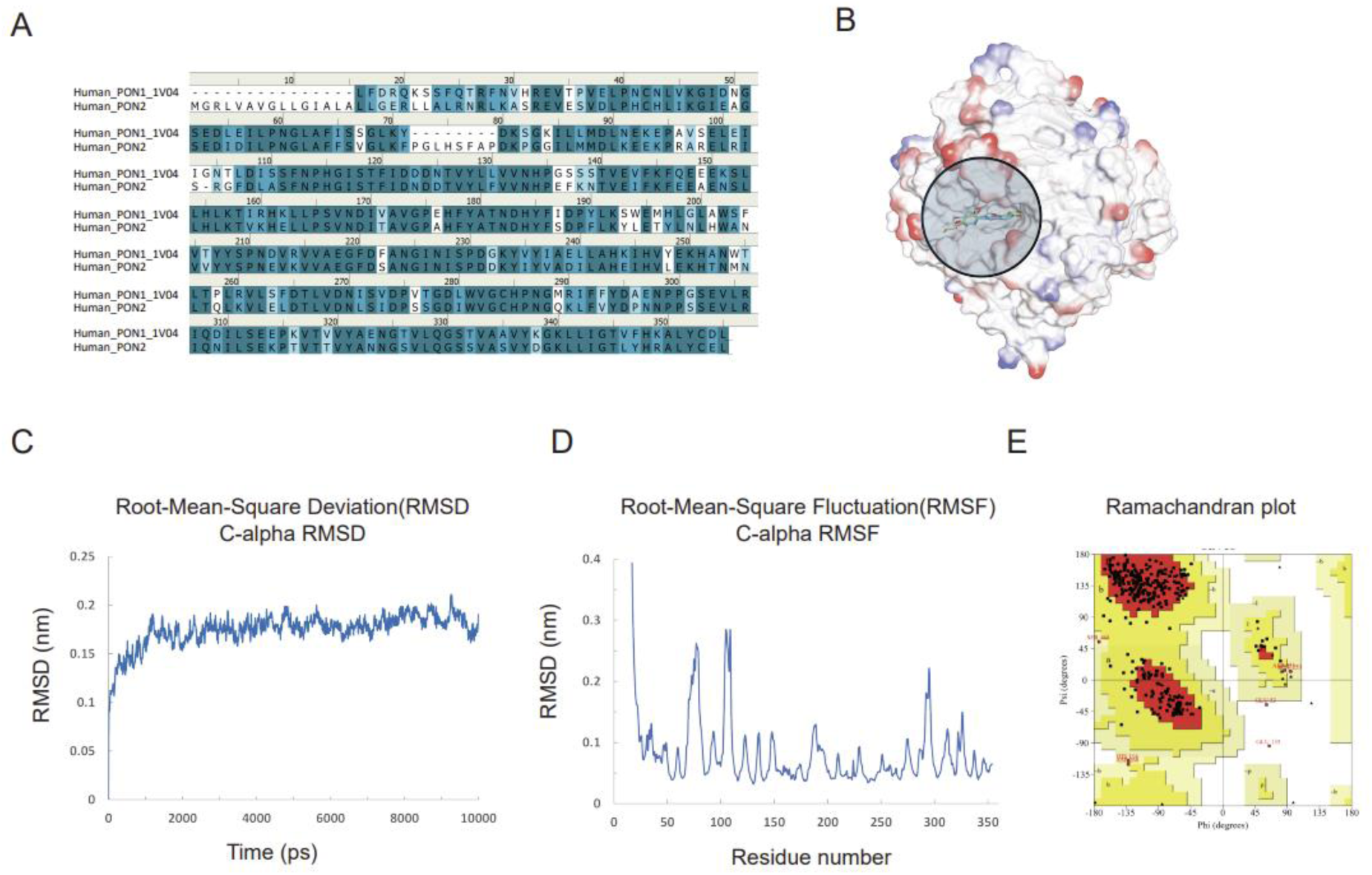
3D structure of PON2 and binding interaction with vutiglabridin. The 3D structure of PON2 was constructed via homology modeling to PON1 and binding interaction with vutiglabridin was assessed. (A) Sequence comparison of PON1 and PON2 proteins show sequence identity of 61.7% and sequence similarity of 79.2%. (B) De novo 3D protein structure of PON2 and the docking site for the tested compounds. (C, D) Molecular dynamic simulation showing RMSD (C) and RMSF (D) graphs for 10 ns show stable maintenance and no significant movement, except for the amino acid residues forming the loop with the end of the N-terminus. (E) Ramachandran plot showing phi(Φ) and psi(Ψ) torsion angles and interatomic collisions. 86.6% of torsion angles were located in the normal distribution area and most amino acids are within the acceptable range.

**Table S1.**
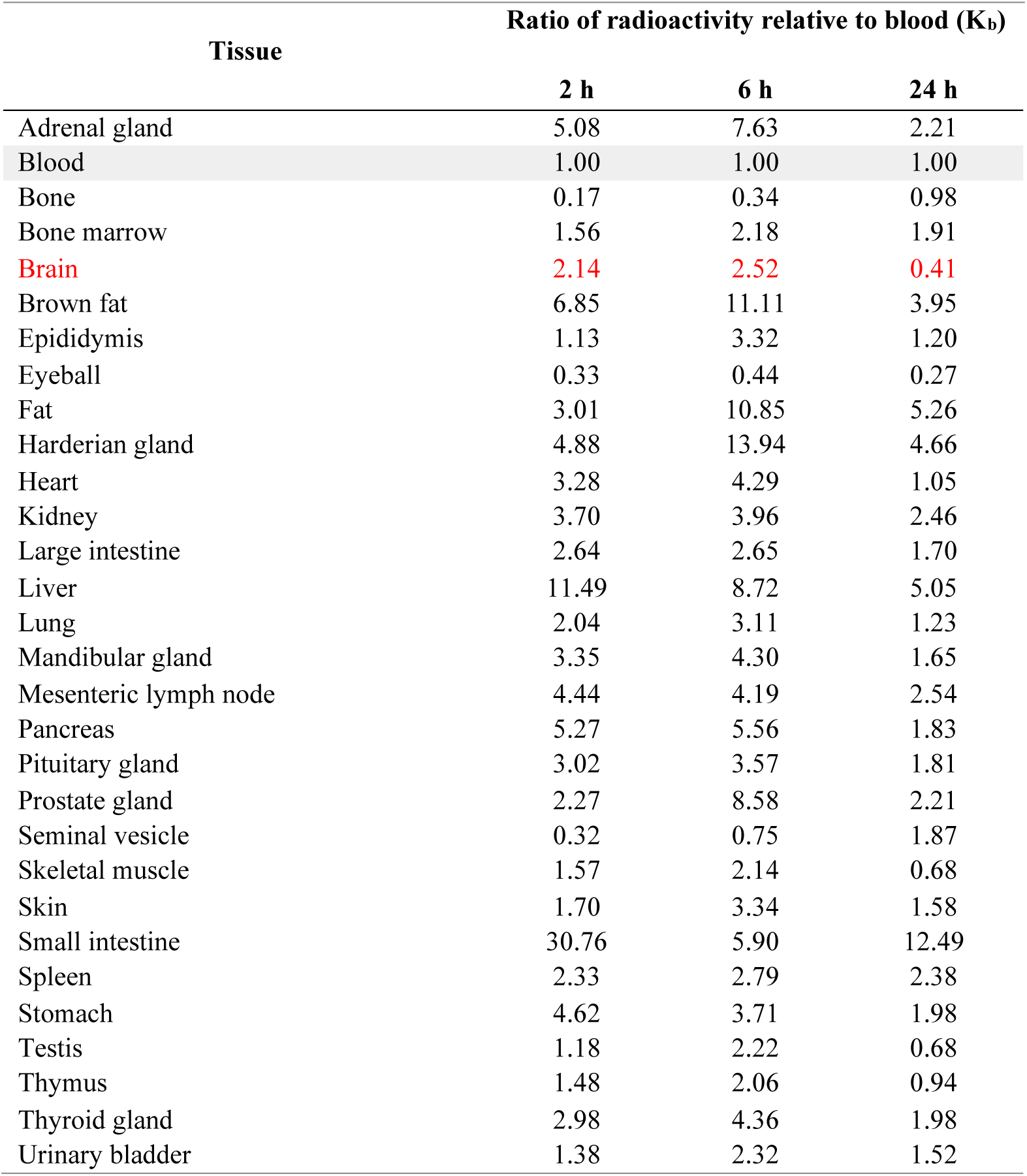
Distribution of radioactive-labelled vutiglabridin in Sprague-Dawley rat. Male Sprague Dawley rats of 8 weeks of age were orally administered 10 mg/kg of ^14^C-labeled vutiglabridin and its distribution throughout the whole body was quantified by measuring radioactivity on the tissues of the whole-body sagittal sections made at 2, 6, and 24 h after the drug administration, and was normalized to the radioactivity in blood (K_b_).

| Tissue | Ratio of radioactivity relative to blood ($K_b$) | | |
| --- | --- | --- | --- |
|  | 2 h | 6 h | 24 h |
| Adrenal gland | 5.08 | 7.63 | 2.21 |
| Blood | 1.00 | 1.00 | 1.00 |
| Bone | 0.17 | 0.34 | 0.98 |
| Bone marrow | 1.56 | 2.18 | 1.91 |
| Brain | 2.14 | 2.52 | 0.41 |
| Brown fat | 6.85 | 11.11 | 3.95 |
| Epididymis | 1.13 | 3.32 | 1.20 |
| Eyeball | 0.33 | 0.44 | 0.27 |
| Fat | 3.01 | 10.85 | 5.26 |
| Harderian gland | 4.88 | 13.94 | 4.66 |
| Heart | 3.28 | 4.29 | 1.05 |
| Kidney | 3.70 | 3.96 | 2.46 |
| Large intestine | 2.64 | 2.65 | 1.70 |
| Liver | 11.49 | 8.72 | 5.05 |
| Lung | 2.04 | 3.11 | 1.23 |
| Mandibular gland | 3.35 | 4.30 | 1.65 |
| Mesenteric lymph node | 4.44 | 4.19 | 2.54 |
| Pancreas | 5.27 | 5.56 | 1.83 |
| Pituitary gland | 3.02 | 3.57 | 1.81 |
| Prostate gland | 2.27 | 8.58 | 2.21 |
| Seminal vesicle | 0.32 | 0.75 | 1.87 |
| Skeletal muscle | 1.57 | 2.14 | 0.68 |
| Skin | 1.70 | 3.34 | 1.58 |
| Small intestine | 30.76 | 5.90 | 12.49 |
| Spleen | 2.33 | 2.79 | 2.38 |
| Stomach | 4.62 | 3.71 | 1.98 |
| Testis | 1.18 | 2.22 | 0.68 |
| Thymus | 1.48 | 2.06 | 0.94 |
| Thyroid gland | 2.98 | 4.36 | 1.98 |
| Urinary bladder | 1.38 | 2.32 | 1.52 |

**Table S2.**
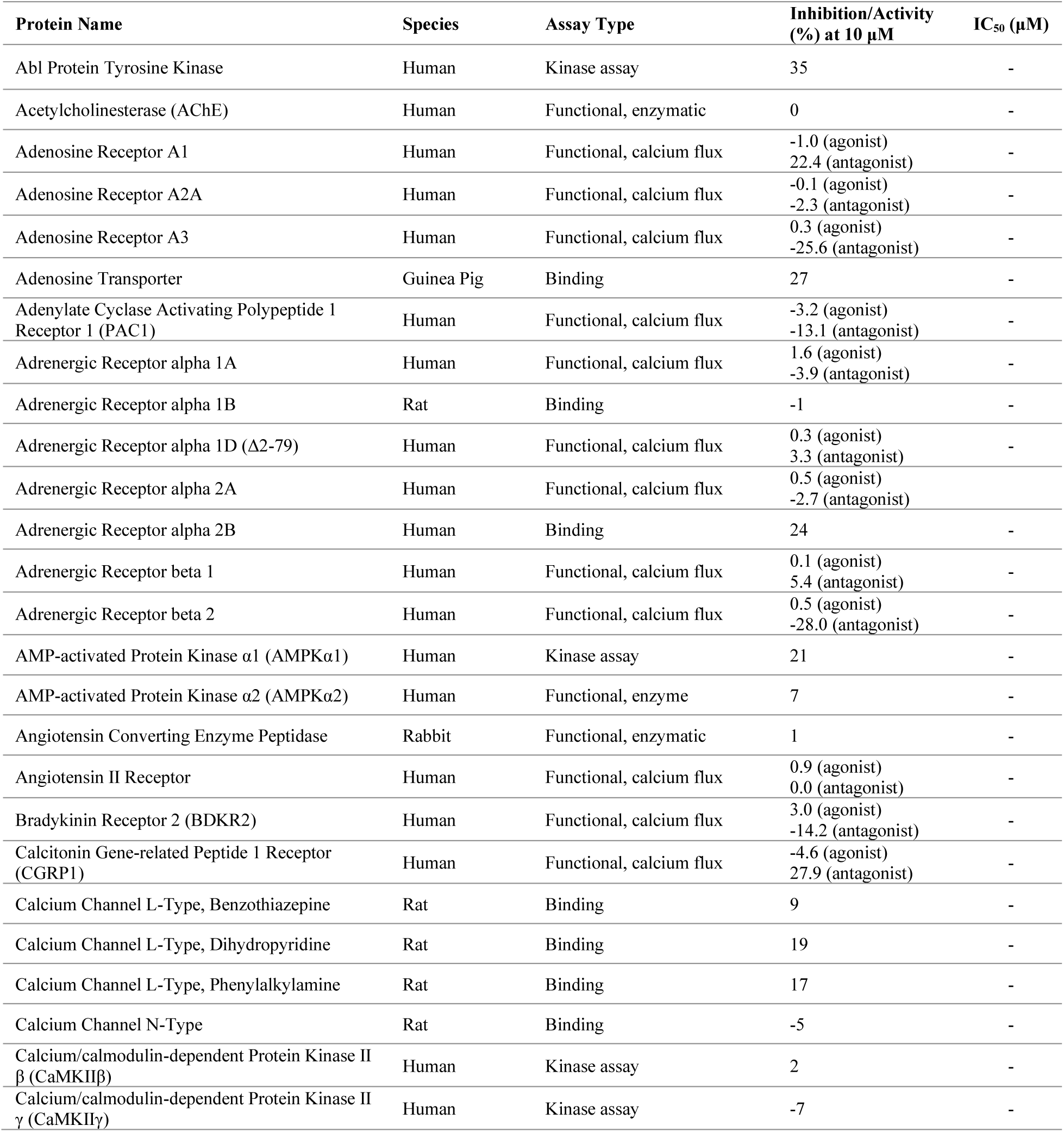
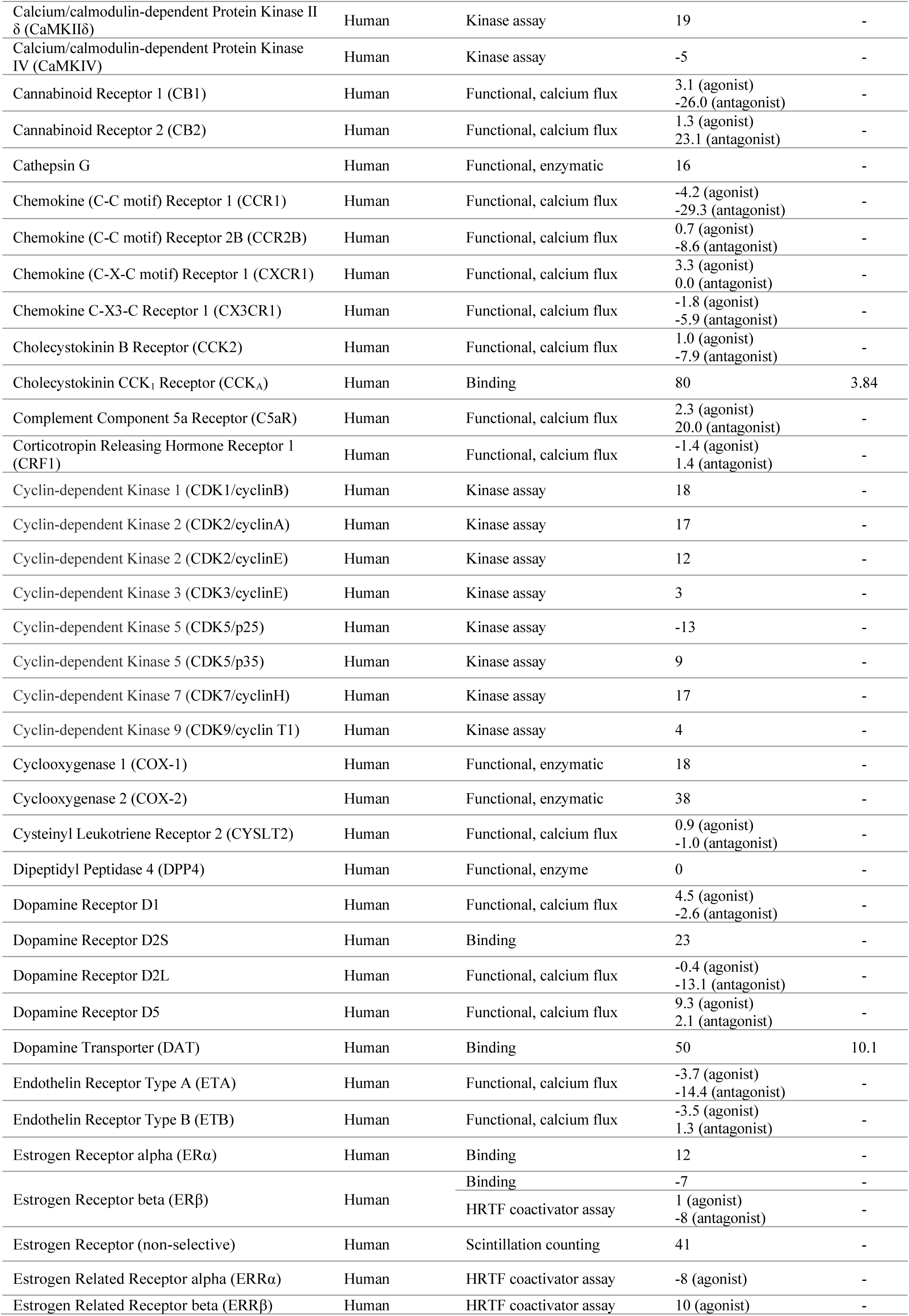
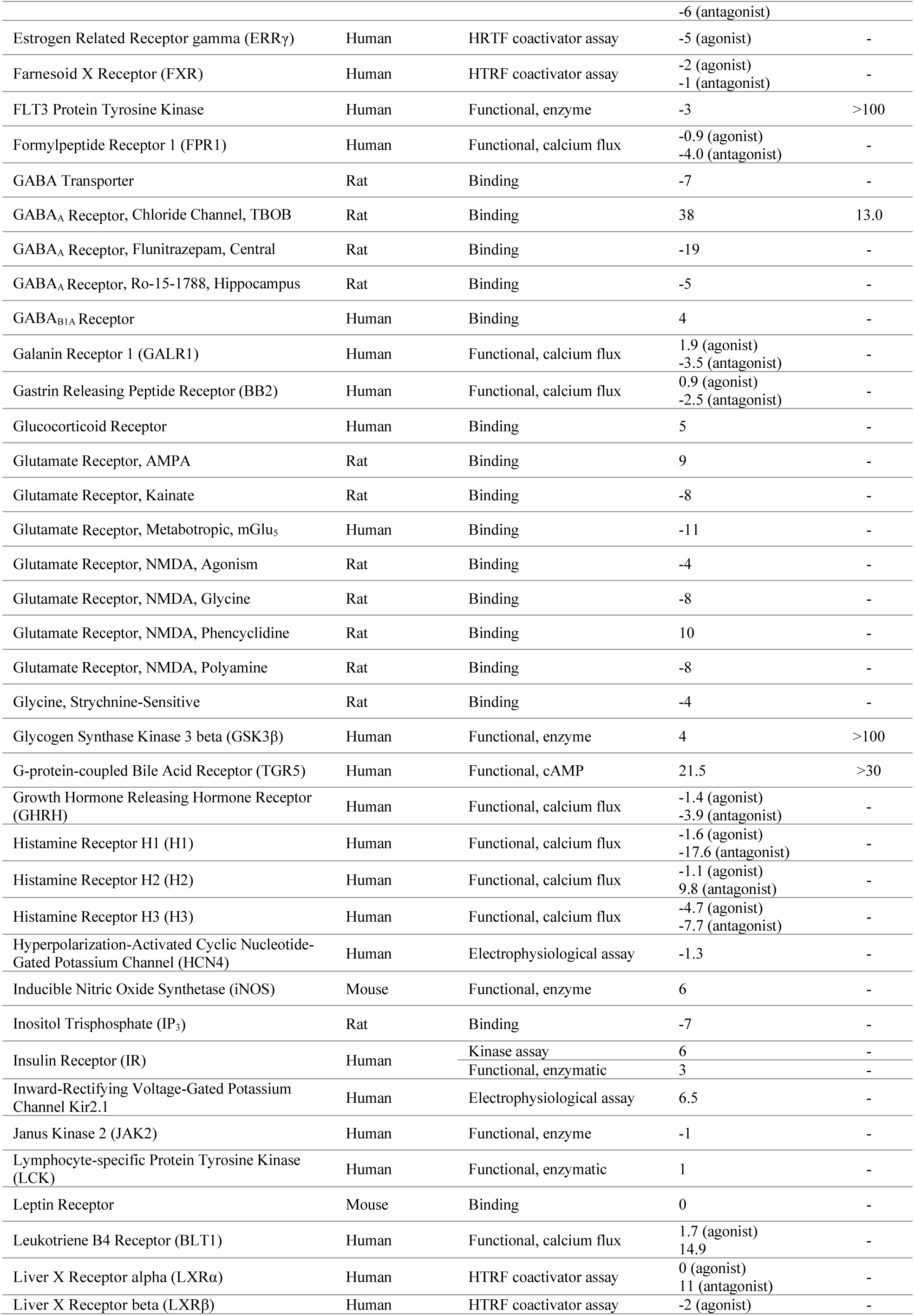
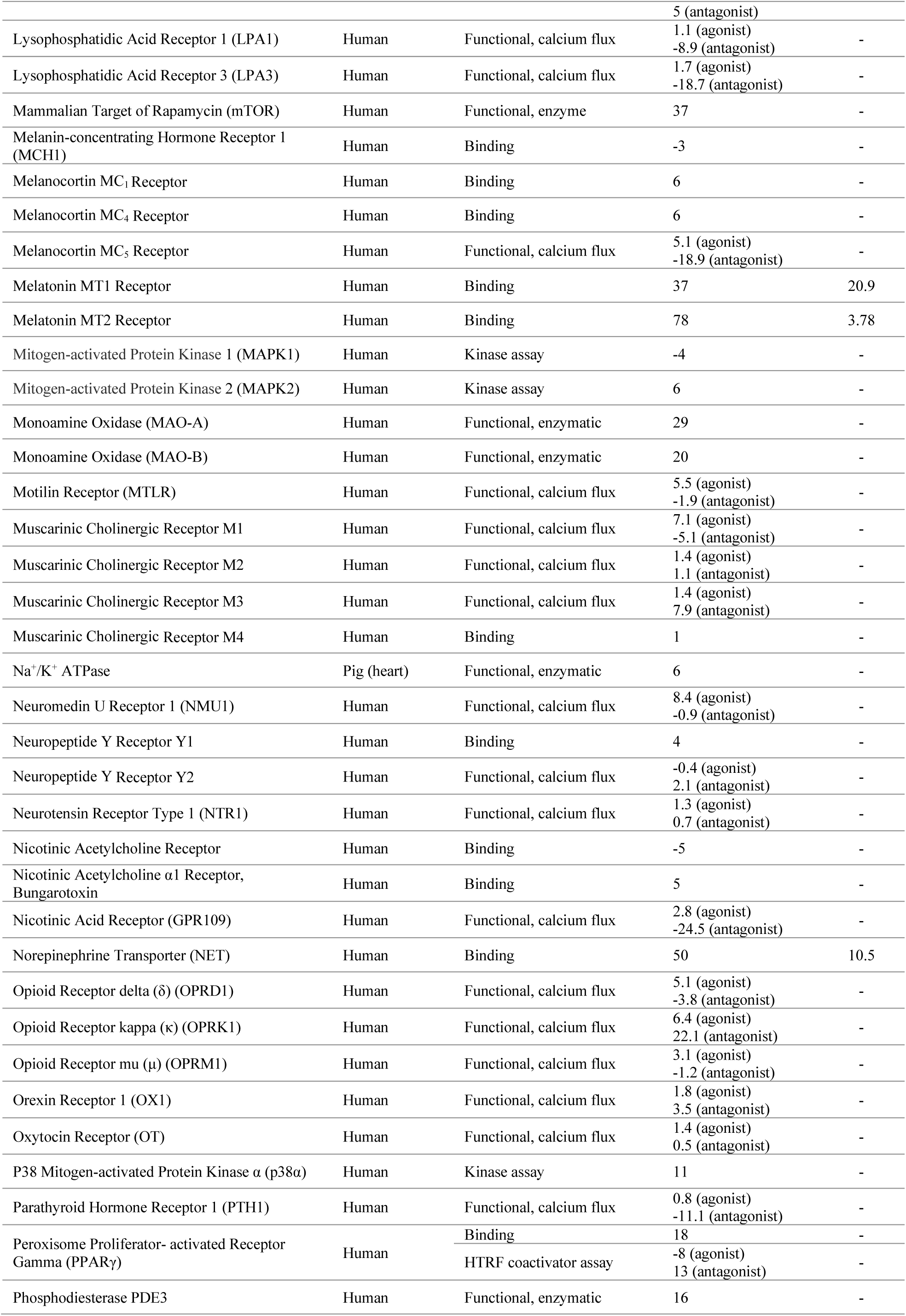
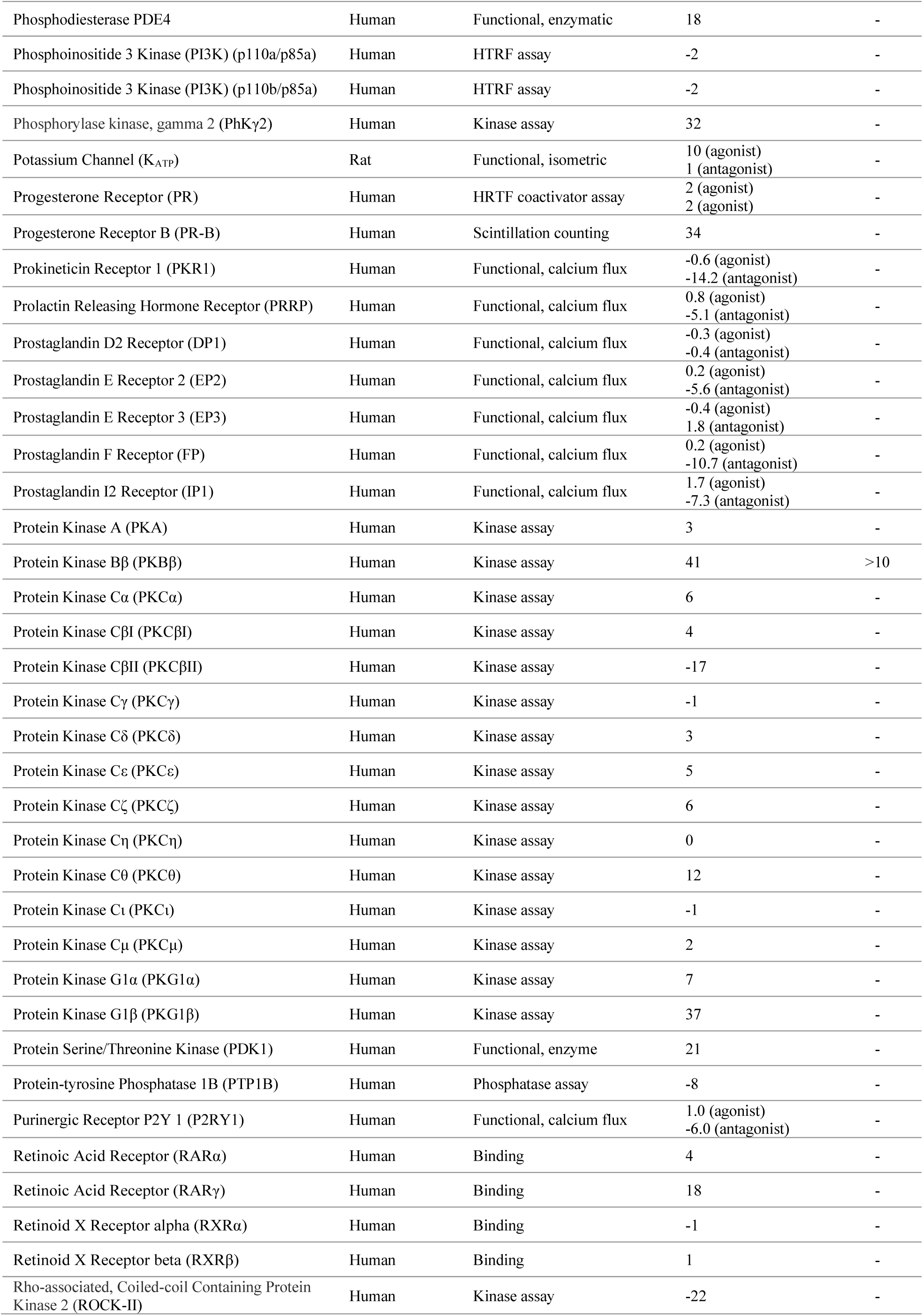
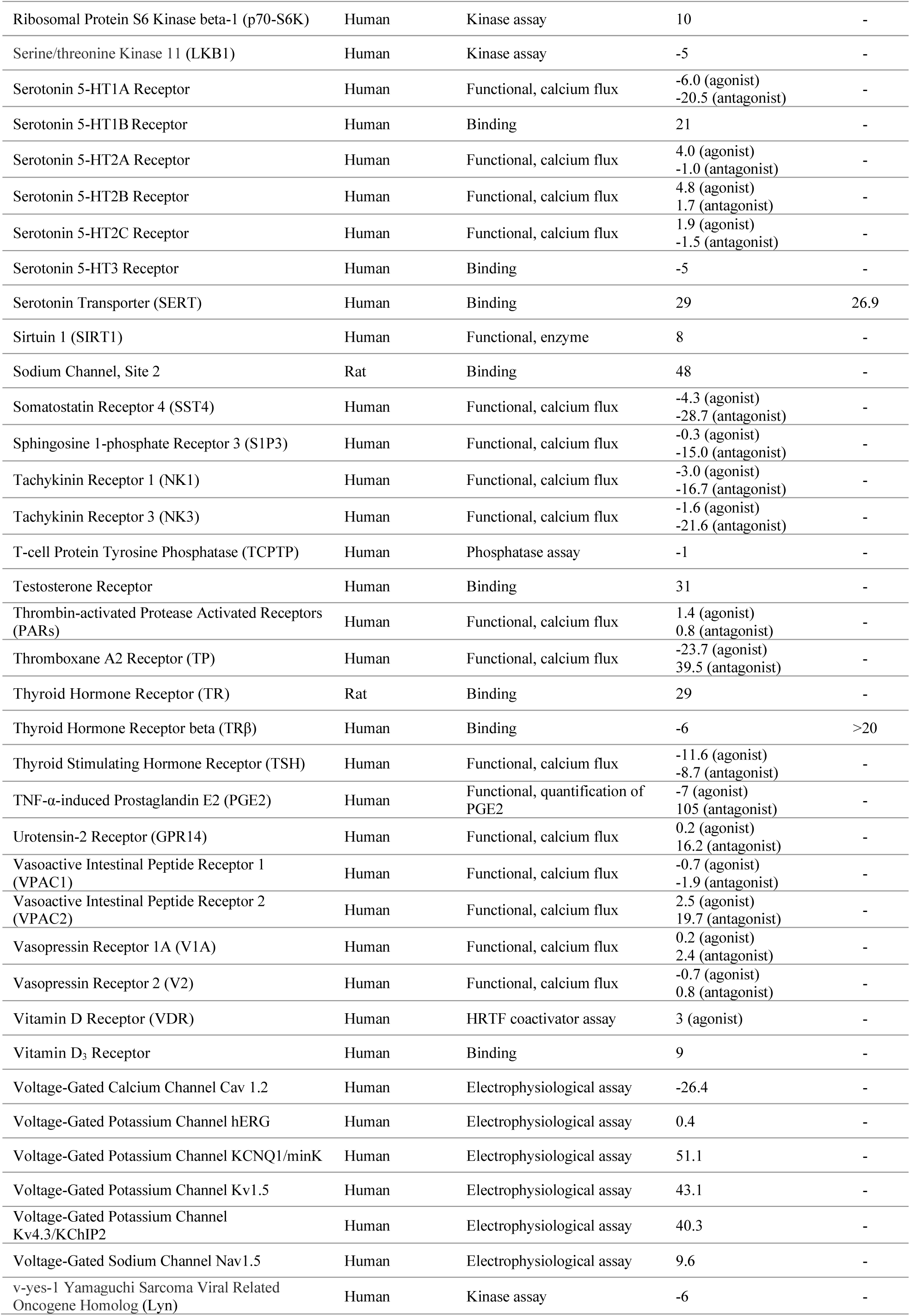
In vitro safety profiling and off-target screening. Assays were performed in accordance with the standard binding, enzymatic, or electrophysical assays established for each of the 214 proteins that include ion channels, GPCRs, nuclear receptors, drug transporters, kinases, and potential targets of vutiglabridin. Inhibition or activity over 50% at 10 μM was used as a cutoff value to proceed with additional kinetic assays to establish IC_50_ values.

**Table S3.**
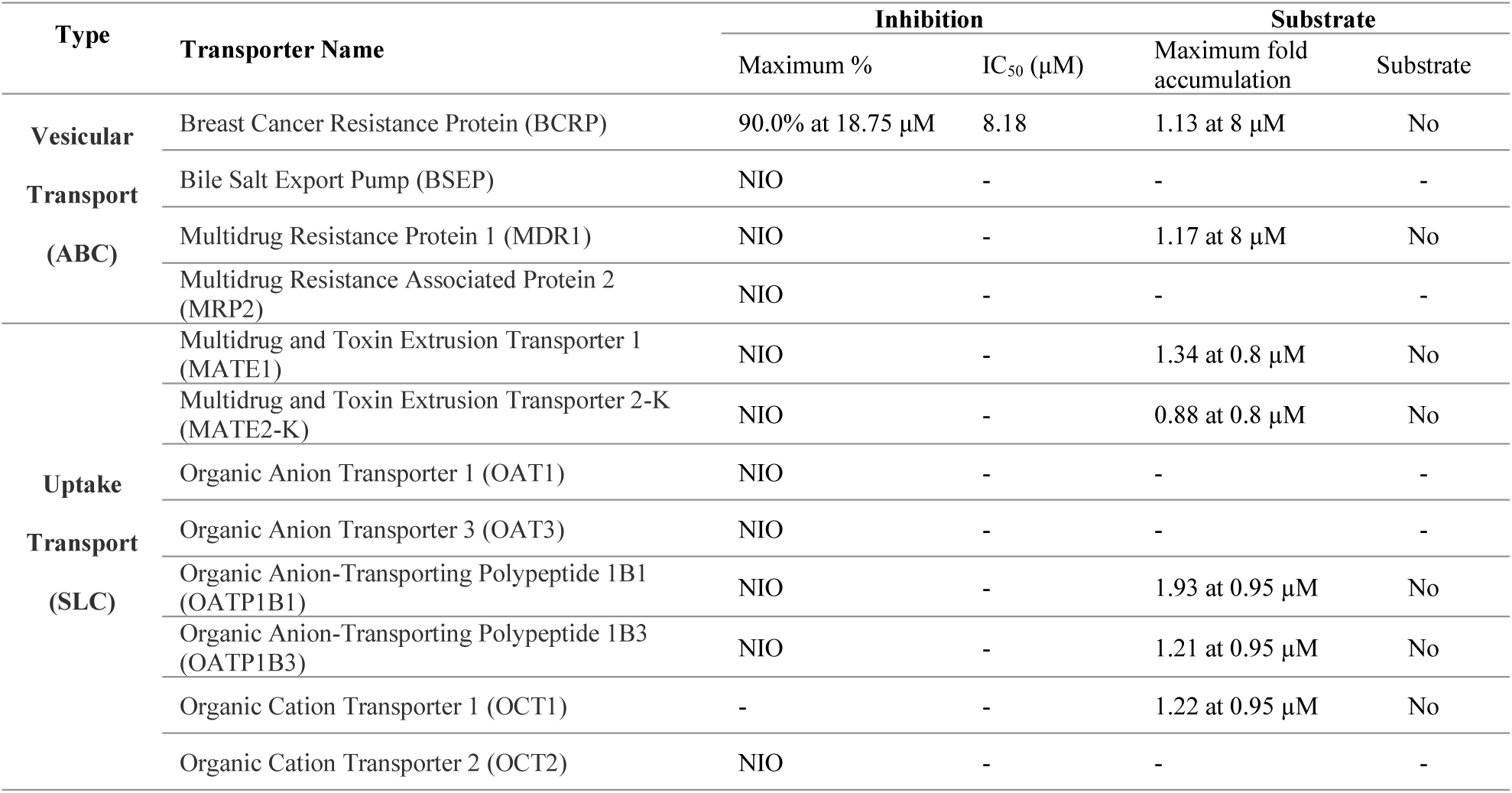
Transporter assay. Vesicular transport and uptake transport inhibition and substrate assays were performed in accordance with the standard assays for each relevant ones of the 12 transporters, respectively. Inhibition under 20% was denoted as NIO (no interaction observed).

**Table S4.**
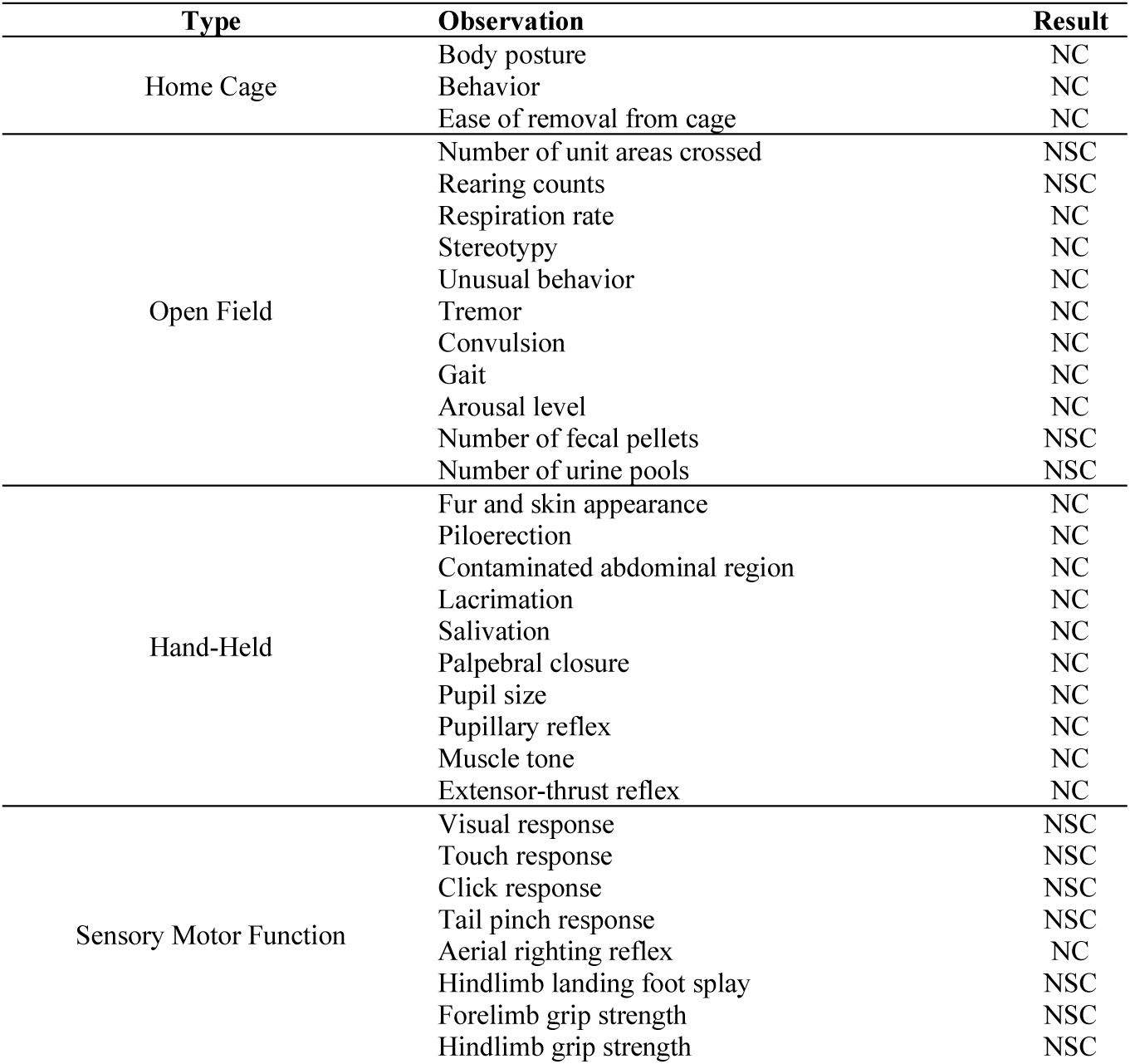
Functional observation battery evaluation. 6-week-old male Sprague-Dawley Rat were orally administered with 0, 500, 1000, or 2000 mg/kg of vutiglabridin (n = 8 per group). Two observers performed blind observations in accordance with the standard guideline (46) at 0.5, 1, 3, 6 and 24 hours post-dose on the basis of the pharmacokinetics parameters. No statistically significant or biologically relevant differences in all parameters were observed for all dose groups at all timepoints. NC = No change, NSC = No statistically significant change.

